# Guild and Niche Determination Enable Targeted Alteration of the Microbiome

**DOI:** 10.1101/2023.05.11.540389

**Authors:** Oriane Moyne, Mahmoud Al-Bassam, Chloe Lieng, Deepan Thiruppathy, Grant J. Norton, Manish Kumar, Eli Haddad, Livia S. Zaramela, Karsten Zengler

## Abstract

Microbiome science has greatly contributed to our understanding of microbial life and its essential roles for the environment and human health^1–5^. However, the nature of microbial interactions and how microbial communities respond to perturbations remains poorly understood, resulting in an often descriptive and correlation-based approach to microbiome research^6–8^. Achieving causal and predictive microbiome science would require direct functional measurements in complex communities to better understand the metabolic role of each member and its interactions with others. In this study we present a new approach that integrates transcription and translation measurements to predict competition and substrate preferences within microbial communities, consequently enabling the selective manipulation of the microbiome. By performing metatranscriptomic (metaRNA-Seq) and metatranslatomic (metaRibo-Seq) analysis in complex samples, we classified microbes into functional groups (i.e. guilds) and demonstrated that members of the same guild are competitors. Furthermore, we predicted preferred substrates based on importer proteins, which specifically benefited selected microbes in the community (i.e. their niche) and simultaneously impaired their competitors. We demonstrated the scalability of microbial guild and niche determination to natural samples and its ability to successfully manipulate microorganisms in complex microbiomes. Thus, the approach enhances the design of pre- and probiotic interventions to selectively alter members within microbial communities, advances our understanding of microbial interactions, and paves the way for establishing causality in microbiome science.

## Main

Microbiome science has contributed greatly to our understanding of microbial life and provided crucial insights on the pivotal roles and capabilities of microbial communities on our planet, from global elements cycling to human health^9–11^. However, we still lack a comprehensive understanding of how these communities are assembled, maintained, and function as a system^6–8^. In particular, the underlying mechanism of microbe-microbe interactions and how microbial communities respond to perturbations remains poorly understood. Current strategies to unravel interactions in microbiomes often include multiple pairwise comparisons of isolates^12–14^ but these studies frequently do not account for higher-order interactions, crucial for understanding and potentially altering heterogeneous communities^15^. Consequently, microbiome science has been largely descriptive and correlation-based, instead of providing accurate predictions centered around mechanistic understanding and established causality^6, 7^. In order to achieve predictive microbiome science we need to comprehensively elucidate the metabolic role of each microbe and its interactions with others. Such knowledge would lead to approaches that rationally change a microbe’s trajectory within a community, for example by selectively promoting or inhibiting its growth.

Here, we present a new technology that integrates transcription and translation measurements to reveal how each microbe allocates its resources for optimal proteome efficiency. mRNA translation into protein is the most energy-demanding process in a cell^16^ and thus microbes closely regulate their resource allocation by prioritizing essential functions through differential translational efficiency (TE)^17, 18^. We hypothesized that direct measurement of TE in a microbial community sample will shine light on the metabolic role of each member of that community and provide a detailed understanding of interactions with other members. We performed metatranscriptomic (metaRNA-Seq) and metatranslatomic (metaRibo-Seq) analysis to directly measure TE *in vitro* in a 16-member synthetic community (SynCom) compiled from rhizosphere isolates grown in complex medium^19^. This approach allowed us to perform a guild-based microbiome classification, grouping microbes according to the metabolic pathways they prioritize, independently of their taxonomic relationship. We demonstrated that guilds predicted competition between members of the same guild with 100% sensitivity and 74% specificity (77% accuracy) in the SynCom. Furthermore, gene-level analysis of import proteins with high TE predicted each microbe’s substrate preferences, i.e. their niche in the community. Such Microbial Niche Determination (MiND) predicted which particular microorganisms would benefit from substrate supplementation with 57% sensitivity and 82% specificity (78% accuracy) in the SynCom. As microbes adapt their translational regulation to community settings, those accurate predictions were not feasible using axenic culture approaches, such as phenotypic microarrays or growth curves. Measurements with limited functional resolution, such as metagenomics or metatranscriptomics alone, did not recapitulate findings obtained by MiND. Combining TE-based MiND and guild predictions allowed us to selectively manipulate the SynCom by increasing or decreasing the relative abundance of targeted members either by adding preferred substrates or by giving an advantage to their competitors. Importantly, the method is scalable to natural samples and can be performed in complex matrices or culture media. We applied MiND and guild classification to native soil and human gut microbiome samples and demonstrated its applicability to forecast changes and alter specific microorganisms in complex microbiomes with high accuracy.

### Guild-Based Microbiome Classification

Bacteria use translational regulation to allocate finite resources and prioritize functions essential for their adaptation^18, 20–22^. Ribosome profiling (Ribo-Seq), i.e. translatomics, allows the direct measurement of protein translation *in vivo* in real time^17, 23^. We have shown previously that translational efficiency (TE), calculated by analyzing the number of ribosomes on a given transcript as the ratio between translated mRNA over total mRNA (Ribo-Seq/RNA-Seq) can be used as a direct readout of functional prioritization in axenic bacterial cultures^18^.

Here, we applied metagenomics, metatranscriptomics, and metatranslatomics^24^ to simultaneously measure TE of multiple organisms in a 16-member microbial community from rhizosphere isolates grown in complex media (see methods, Fig. 1a). These multi-omics data showed excellent reproducibility between biological replicates and highlighted substantial differences between metagenomic, -transcriptomic, and -translatomic data (Suppl. Fig. S1).

**Fig. 1.**
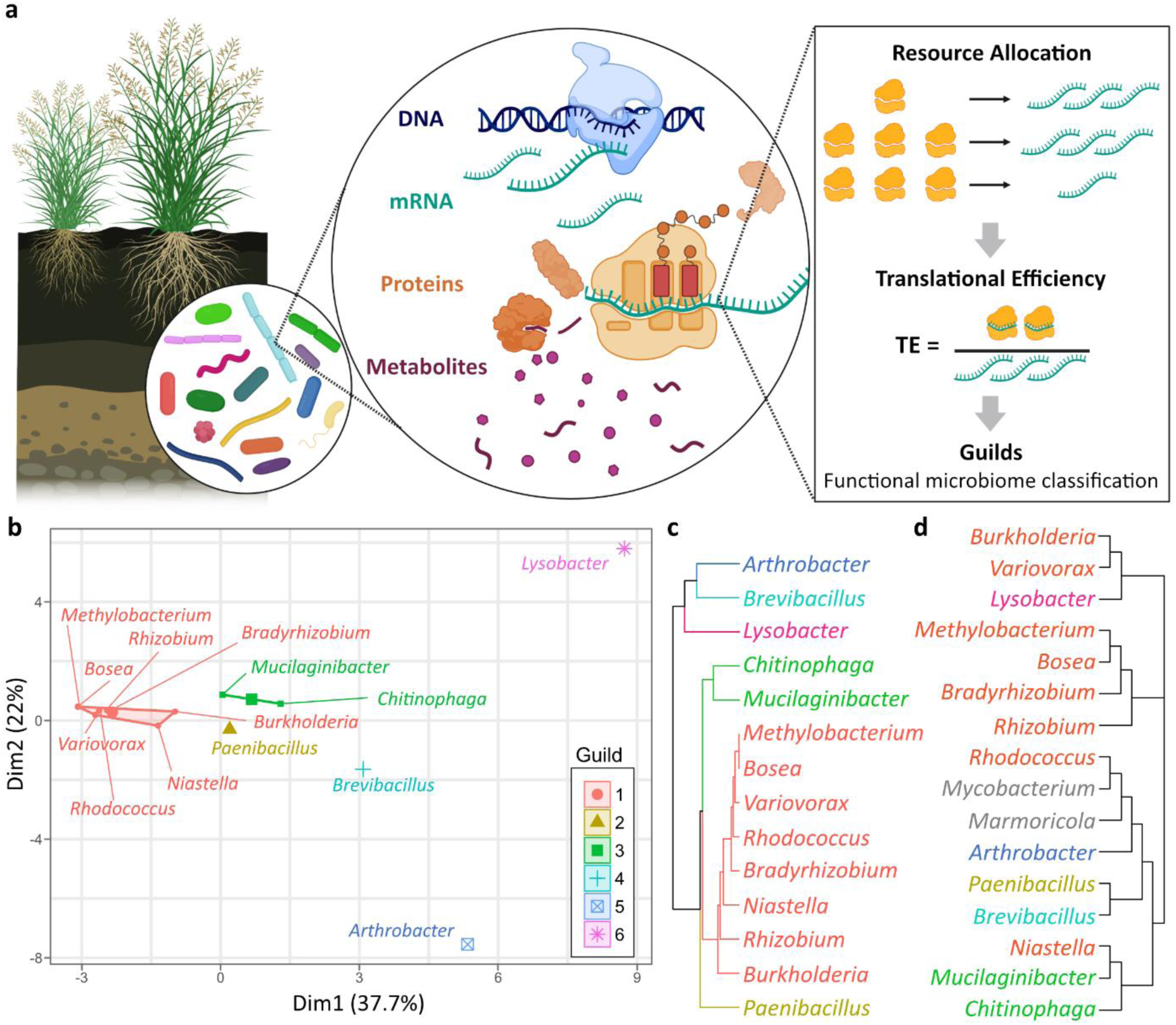
Guild-based microbiome classification of a 16-member SynCom based on translational efficiency (TE). **a)** Conceptual overview of TE as a readout of functional prioritization for each microbe in a 16-member SynCom isolated from the switchgrass rhizosphere, where each member has a limited amount of resources (ribosomes) to allocate for protein translation. TE is computed as the metaRibo-Seq/metaRNA-Seq ratio, i.e. the ratio between translated mRNA and total mRNA detected at one given instant in the sample; **b)** PCA cluster plot and **c)** dendrogram of TE on 275 KEGG^25^ metabolic pathways (n=4 replicates) allowed classification of the 16 SynCom members into 6 different guilds, in which microbes share similar metabolic pathway prioritizations, as detailed in Suppl. Table S1 and in Suppl. Fig. S2; **d)** phylogenetic tree based on 16S rRNA sequences shows substantial differences with the TE-based guild dendrogram **c)**, indicating that guilds are not based solely on phylogeny. Note, two bacteria (*Mycobacterium* and *Marmoricola*) had low KEGG pathway coverage due to low abundance in the community (0.04 and 0.13 % of total reads, respectively), and thus are absent from **b)** and **c)**.

We categorized the SynCom members into functional groups or guilds, based on the metabolic pathways they prioritized (i.e. TE profiles) (see methods, Fig. 1b,c). The 16-member SynCom was divided into 6 guilds, defined by specific metabolic functions (i.e. pathway prioritization) (Fig. 1b,c). For example, *Lysobacter* (guild 6) has a significantly higher TE for denitrification and dissimilatory nitrate reduction compared to other guilds, while *Chitinophaga* and *Mucilaginibacter* (guild 3) comprise a high TE for assimilatory sulfate reduction, thiosulfate oxidation, and multiple antimicrobial resistance pathways (Suppl. Table S1, Suppl. Fig. S2).

Metabolic pathway prioritization differed between bacteria grown axenically or in the SynCom, highlighting the importance of performing functional analysis directly in community settings (Suppl. Fig. S3). The TE-based guilds were substantially different from phylogenetic clustering, indicating that functional categories often are independent of taxonomic relationship (Fig. 1d, Suppl. Fig. S4). Guilds were also dissimilar to cluster information obtained from genome content, metatranscriptomics, or metatranslatomics data alone (Suppl. Fig S4). Combined, this data hints at current limitations of 16S rRNA and genome-based approaches that infer function and activity from phylogeny or genome content alone.

### Guilds predict bacterial competition

Ecological guilds are defined as functional categories molded by adaptation to the same class of resources but also by competition between its members^26, 27^. To test if such within-guild competition applies to microbes, we experimentally removed individual members from the SynCom and evaluated the effect on relative abundance of the remaining 15 members (Fig. 2e).

**Fig. 2.**
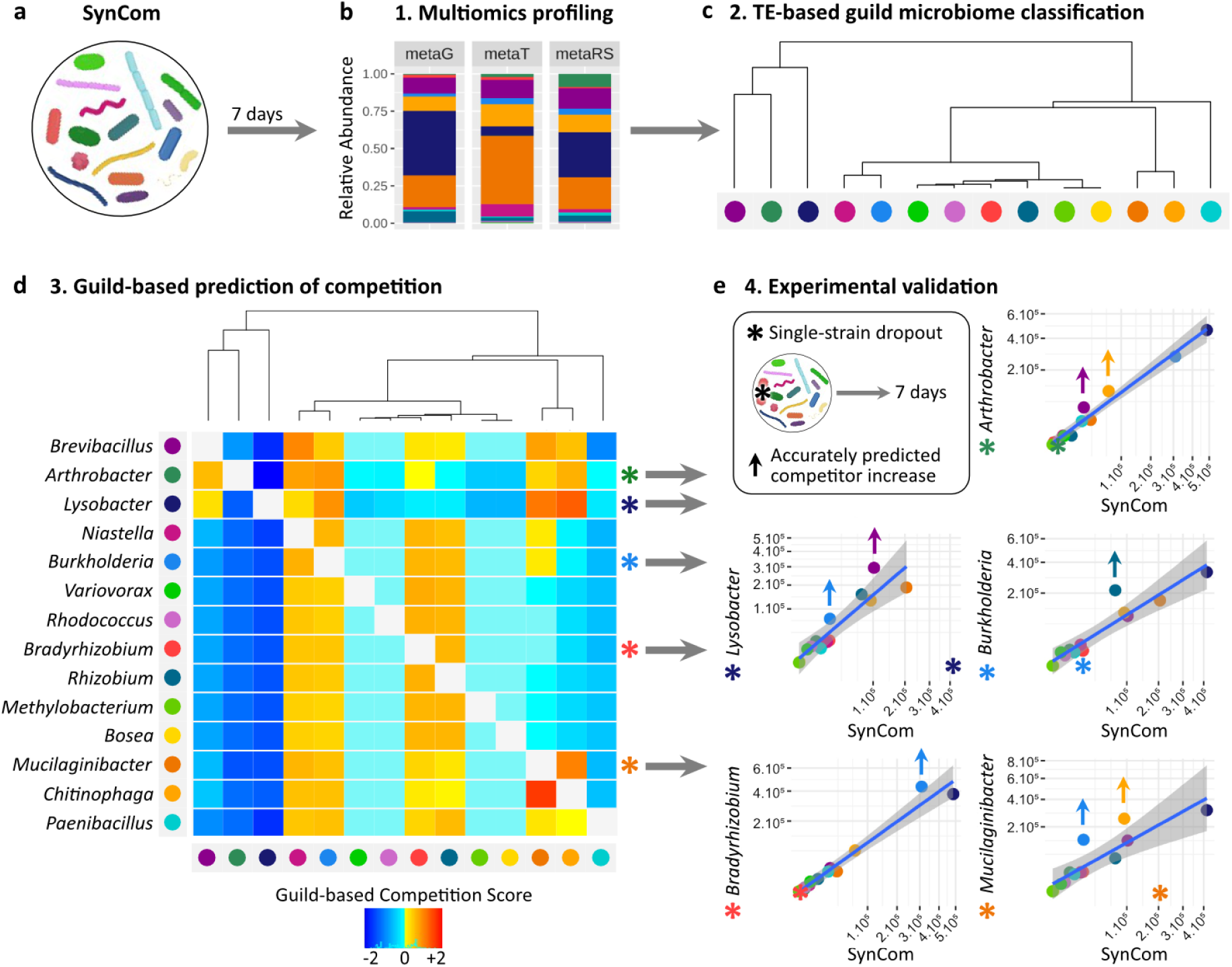
Guild-based microbiome classification predicts competition interactions in a microbial community. a-b) We performed multi-omic profiling of a 16-member soil SynCom; **b)** relative abundances of the 16 SynCom members at the metagenomic, metatranscriptomic and metatranslatomic levels (average of replicates shown in Suppl. Fig. S1 a), color key d) applies; **c)** metatranslatomic (metaRS) and metatranscriptomic (metaT) profiles were used to compute TE and to classify members into guilds as described in Fig. 1b,c; **d)** we computed a competition score to predict competitive interactions against each SynCom member based on the proximity of its guild with each other member’s (see methods). High competition scores (warm colors) indicate SynCom members (in columns) that are likely to compete with the targeted member (in row). Asterisks indicate competition against targeted members that have been tested experimentally as shown in e); **e)** five individual members were experimentally removed from the SynCom prior to incubation (asterisks), and relative abundances of all remaining members were compared to the non-modified SynCom. Graphics show linear regression and 99% confidence interval (CI, in gray) of square-transformed relative abundances (RPKM) in the SynCom (x-axis) vs. single dropout SynCom (y-axis). Organisms above the 99% CI (arrows) were considered significantly increased in response to the dropout, thus showing that they were competing with the removed member in the SynCom. In all the tested conditions, significantly increased members were accurately predicted to compete with the dropped out member (high competition scores against the targeted member in d, rowwise).

Fig. 2b shows the relative abundance of each of the 16 SynCom members at the metagenomic, metatranscriptomic, and metatranslatomic level (Fig. 2a,b). We computed a guild-based competition score reflecting the guild clustering distance matrix, with the hypothesis that similar guilds would predict competitive interactions (see methods, Fig. 2c,d, Suppl. Fig S5). In 5/5 tested conditions, a single microbe dropout benefitted at least one of its closest competitors from the same guild whose relative abundance increased significantly (Fig. 2d,e). For example, when *Mucilaginibacter* was removed we observed that two of its closest neighbors in the guild clustering (*Chitinophaga* and *Burkholderia*) increased in abundance (Fig. 2e, bottom right). Similarly, removal of *Burkholderia* resulted in an increase of its close competitor *Rhizobium* (Fig. 2e, middle right). Comparable results were obtained for all single member dropout experiments, i.e. removal of *Arthrobacter*, *Bradyrhizobium*, *Burkholderia*, *Lysobacter*, and *Mucilaginibacter* (Fig. 2e). We also observed elevated metabolic activity based on metatranscriptomic and metatranslatomics levels for microbes that increased in abundance (Supp Fig. S6). Overall, the guild-based competition scores allowed us to predict competitive interactions within the microbial community with excellent sensitivity (100%) and specificity (74%).

To further validate the competitive interactions, we conducted an in-depth analysis on two of the strong competition pairs observed in the dropout experiments (i.e. *Chitinophaga*- *Mucilaginibacter* and *Burkholderia*-*Rhizobium*, Fig. 2e, Suppl. Fig. S7 a,e). Complementary to the dropout experiments, we observed that adding more cells of each of these members to the SynCom (similar to a probiotic intervention) resulted in a very specific decrease in relative abundance of their main competitor (Suppl. Fig. S7 b,c,f,g). On the other hand, experimentally removing both *Chitinophaga* and *Mucilaginibacter* or *Burkholderia* and *Rhizobium* had little to no effect on the abundance of microbes from the other guilds (Suppl. Fig. S7 d,h), suggesting that competition is restricted to each guild. We then performed spot-on-lawn assays^28^ to screen for antimicrobial compounds produced by these competition pairs. In line with guild-based predictions of competition, *Chitinophaga* specifically inhibited the growth of *Mucilaginibacter*, while it failed to inhibit growth of any other SynCom member (Suppl. Fig. S8). This hints at the production of narrow-spectrum antimicrobials by *Chitinophaga* specifically targeted against its guild competitor *Mucilaginibacter*. In contrast, *Burkholderia* and *Rhizobium* did not inhibit each other or any other member’s growth, suggesting that the mechanism of competition within this guild is likely not augmented by antimicrobials (Suppl. Fig. S9). Our data confirms that TE-based guilds accurately predict competitive interactions in a microbial community. Of note, clustering of guilds and prediction of competitions based on TE information (100% sensitivity and 74% specificity) outperformed analysis based on metatranscriptomic (37.5% sensitivity and 76% specificity, Suppl. Fig. S10) and was more sensitive than analysis based on metatranslatomic data (75% sensitivity and 81% specificity, Suppl. Fig. S10) alone.

### Microbial Niche Determination (MiND)

Next, we used TE information to identify substrate preferences, i.e. metabolites that would specifically promote growth of selected members of the SynCom, akin to a prebiotic. We hypothesized that high TE for genes coding for import proteins would indicate prioritized metabolism for the corresponding substrates, allowing for Microbial Niche Determination (MiND).

A total of 88 genes coding for import proteins were detected in the SynCom at the metagenomic level, of which 40 genes (45%) were transcribed and translated (Suppl. Fig. S11). We performed MiND by calculating the TE for each of these 40 import protein genes in each SynCom member thus determining their substrate preference, i.e. their niche (Suppl. Fig. S12). Based on this analysis we selected metabolites to be tested as prebiotic interventions in the SynCom with the goal to selectively alter its composition (see methods).

The ability to utilize substrates predicted by MiND was validated for each SynCom member through phenotypic microarrays (Suppl. Table S2, Suppl. Fig. S13) and growth assays (Suppl. Fig. S14). A total of 89% of substrates with high TE importers measured in the SynCom were confirmed as growth supporting in axenic culture. In contrast, the ability to utilize a substrate in isolation did not necessarily translate into a high priority for this substrate’s consumption in the SynCom. Only 39% of substrates metabolized in axenic condition translated into a high TE for this substrate’s import protein(s) in the SynCom (Suppl. Table S3). This highlights that bacteria possess the ability to utilize a range of substrates in axenic culture, but will only prioritize a fraction of those, i.e. their niche, when growing in a community.

### MiND predicts effects of substrate addition

We supplemented the complex culture medium (R2A) with metabolites identified by MiND to benefit microbes that prioritize the import of these substrates (i.e. primary targets). In addition, we hypothesized that metabolite-induced increased abundance of selected bacteria would result in a concomitant decrease of their nearest guild competitors (i.e. secondary targets). A total of 11 compounds were tested in three different concentrations, including six sugars (fructose, galactose, maltose/maltodextrin, ribose, trehalose, xylose), two diamines (putrescine, spermidine), one amino acid (glutamate), one peptide (glutathione), and one inorganic compound (sulfate/thiosulfate) (Suppl. Fig. S15-S16).

As predicted, primary targets were specifically increased in relative abundance upon addition of their preferred metabolite (Fig. 3, Suppl. Fig. S12, S15-S16). At the same time, when primary targets increased, secondary targets (competitors from the same guild) significantly decreased in abundance, with no or non-significant effects on non-competitors. For example, addition of ribose induced a predicted increase of the targets *Paenibacillus* and *Burkholderia* (primary targets), which exhibited the highest TE for ribose importers (RbsA, RbsB, RbsC) in the SynCom; concurrently, *Burkholderia*’s competitors *Variovorax*, *Rhizobium*, and *Bradyrhizobium* (secondary targets) decreased in abundance (Fig. 3d,e,h, Fig. 1b,c, Fig. 2d). Similarly, addition of glutathione increased the primary targets *Burkholderia* and *Chitinophaga*, both having a high TE for the glutathione import protein GsiA, while decreasing their competitors *Mucilaginibacter* and *Bradyrhizobium* (Fig. 3d,f,i, Fig. 1b,c, Fig. 2d). Addition of putrescine increased *Paenibacillus* and *Rhodococcus*, which both had a high TE for putrescine import proteins PuuP and PotA, while reducing the competitors *Rhizobium*, *Bradyrhizobium*, and *Mucilaginibacter* (Fig. 3d,g,j, Fig. 1b,c, Fig. 2d). Overall, MiND predicted the increase of primary target(s) for 9/11 tested substrates, with 57% sensitivity and 82% specificity (78% accuracy) (Suppl. Table S3). In all of these 9 cases, the successful increase of the primary targets also resulted in a decrease of at least one of their competitors (secondary targets). The guild classification predicted such competition-based decrease of secondary targets with 93% sensitivity and 65% specificity (70% accuracy) (Suppl. Fig. S9) (Suppl. Table S3).

**Fig. 3.**
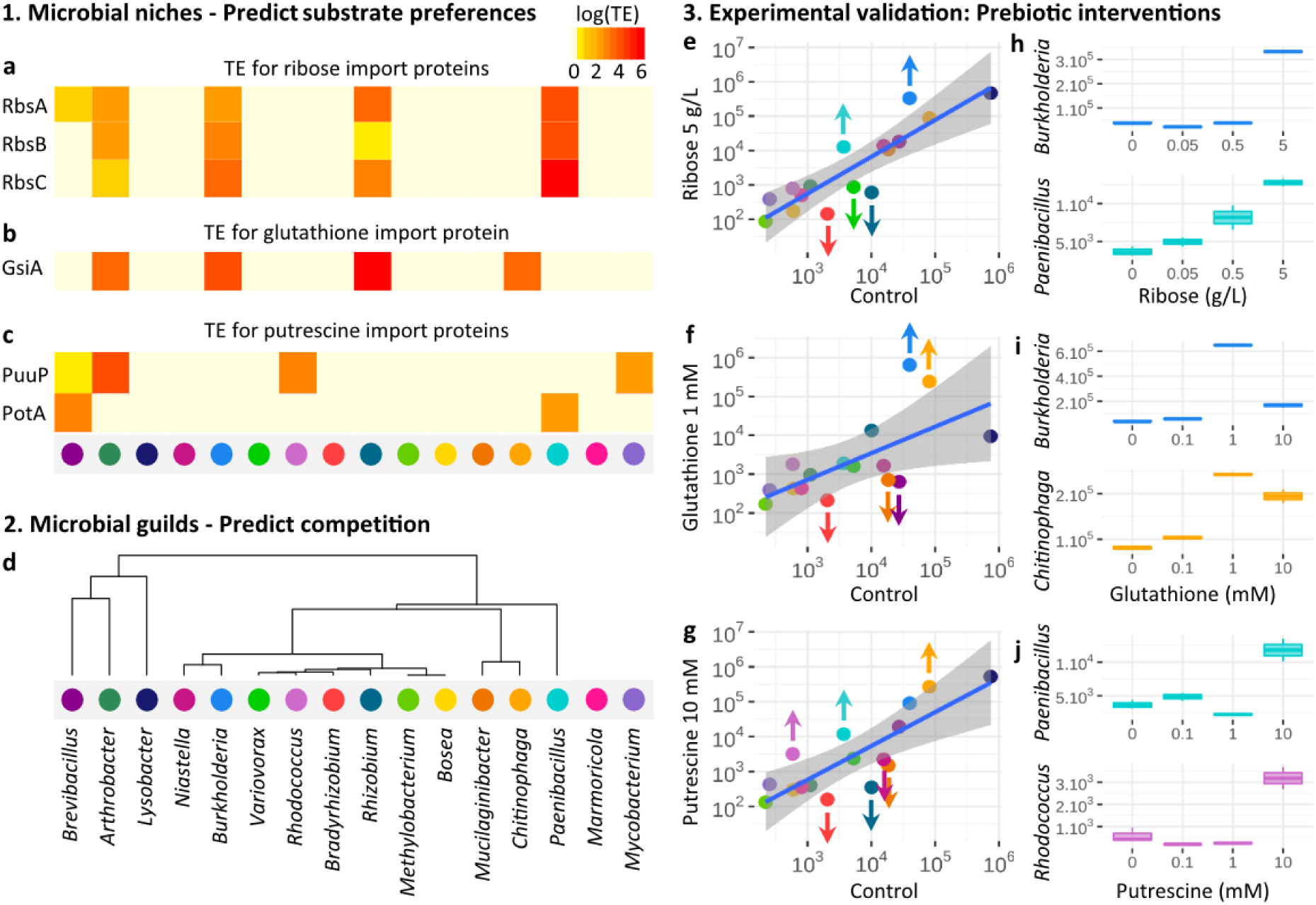
TE-based microbial niches and guilds accurately predict the effect of substrate addition in the SynCom in three steps. a-c) 1. MiND was performed by measuring TE for import proteins in a 16-member SynCom grown in complex medium and was employed to predict each microbe’s substrate preferences. **a**,**b**, and **c** show TE for ribose, glutathione, and putrescine import proteins, respectively. Organisms with high TE for a substrate’s import protein were predicted to increase in relative abundance upon addition of this substrate as a prebiotic to the culture medium (primary targets); **d)** 2. Guilds were used to predict competition in the SynCom as described in Fig. 2c,d. Competitors of the organisms that benefit from a prebiotic intervention were predicted to decrease in relative abundance (secondary targets); **e-j)** 3. Predictions made in a-d) were experimentally validated by supplementing the SynCom culture medium with selected substrates; **e-g)** linear regression and 99% CI of metagenomic relative abundances (RPKM, log scaled) in 0.1x R2A control versus 0.1x R2A + ribose (**e**), glutathione (**f**) or putrescine (**g**). Microorganisms above or below the 99% CI are considered significantly increased or decreased upon metabolite addition (arrows). As predicted, 6/7 organisms that increased in relative abundance had a high TE for the added substrate’s import protein(s), and 9/10 microbes that concomitantly decreased were competitors from a similar guild (see Fig. 2 for detailed competition scores). Note: we did not predict *Brevibacillus* decrease in f) and *Chitinophaga* increase in g); **h-j)** boxplots showing RPKM abundance of significantly increased primary targets with increasing concentration of ribose (**h**), glutathione (**i**), or putrescine (**j**). More examples are available in Suppl. Fig S12, S15 and S16.

Overall, combining MiND and guild classification accurately predicts substrate preferences and competition in a 16-member SynCom and can be employed to design and predict the outcome of targeted interventions in a microbial community (Suppl. Fig. S17).

### MiND predicts intervention outcomes in soil

After benchmarking MiND for the 16-member SynCom, we evaluated if principles derived from guild and niche elucidation could be extrapolated to the soil environment, harboring one of the most complex microbiomes. For this, soil samples from the switchgras*s* (*Panicum virgatum*) rhizosphere, similar to the site from which the SynCom strains were originally isolated^19^, were incubated in 30 different conditions: 0.1x R2A alone (soil control), with and without metabolite additions (i.e. prebiotic, 7 treatments tested), with and without individual SynCom member additions (i.e. probiotic, 8x10^5^ CFU/mL, 7 treatments tested) or the entire 16-member SynCom addition (i.e. probiotic consortium, 8x10^5^ CFU/mL of each member), as well as 14 treatments of combined pre- and probiotic conditions (see methods, Fig. 4a, Suppl. Fig. S18). Our data showed high reproducibility between replicates (Suppl. Fig. S18). Multi- omics data from soil + SynCom samples confirmed that guilds and niches were similar to those observed in the SynCom alone (Fig. 4b, Suppl. Figs. S19, S20, as compared to Fig. 1c, Suppl. Fig. S12), suggesting that guilds and niches are stable under the tested conditions and are independent of the size and diversity of the microbial community.

**Fig. 4.**
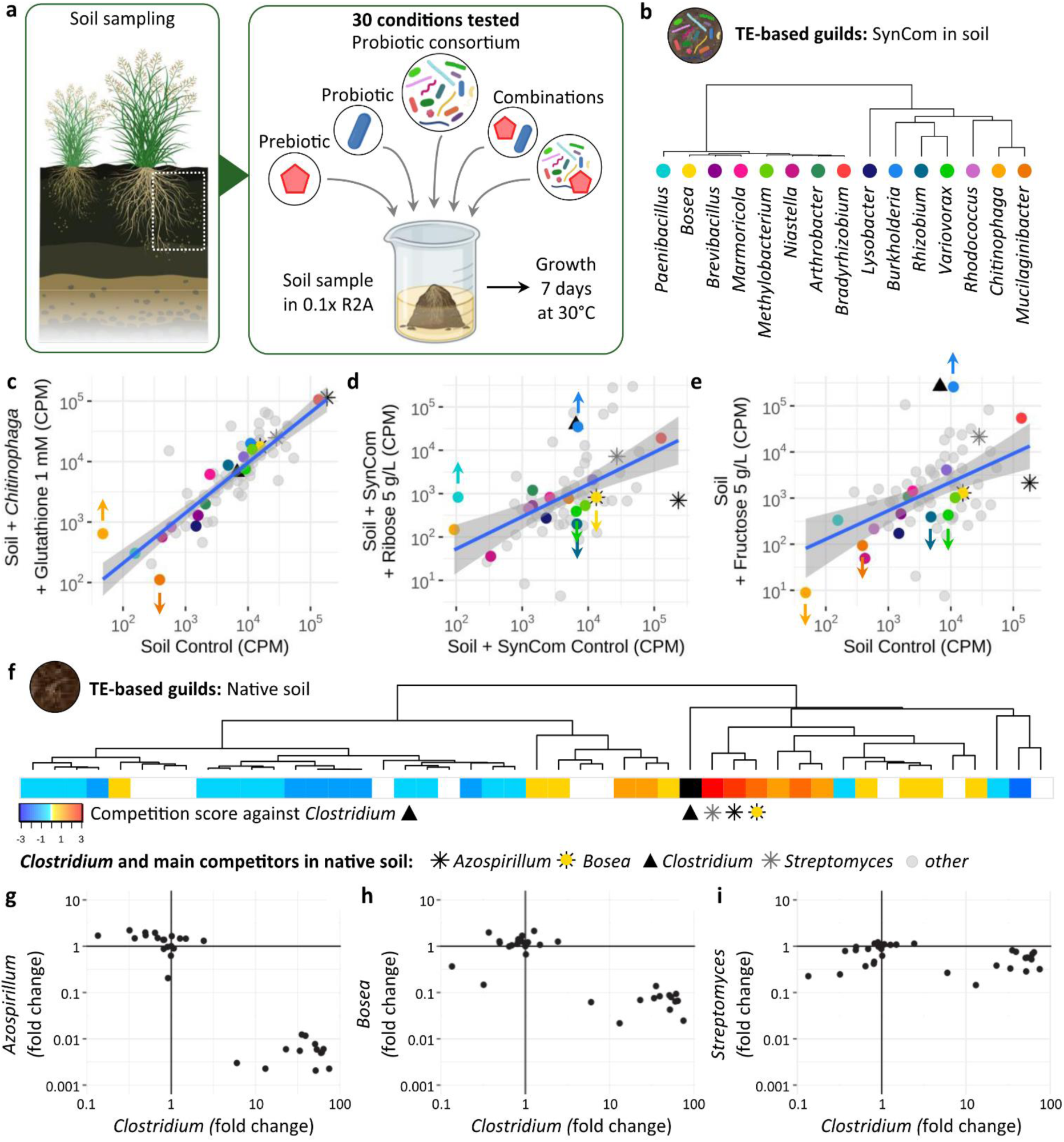
Guilds and MiND accurately predict pre- and probiotic treatment outcomes in soil. **a)** Probiotic and/or prebiotic interventions were carried out by adding a single probiotic, probiotic consortium (i.e. SynCom), and/or prebiotic to the soil prior to growth at 30 °C for 7 days (see methods). Interventions were designed to increase primary targets based on their niche (Suppl. Figs. S12, S19), and decrease secondary targets based on guild competition (see **b**); **b)** TE-based guild clustering to predict competition between SynCom members grown in soil (competition scores are displayed in Suppl. Fig. S20); **c-e)** linear regression and 95% CI of metagenomic relative abundances (CPM, log scaled) in control (x-axis) versus tested pre- and/or probiotic conditions (y-axis); **c)** successful increase of primary target *Chitinophaga* and decrease of secondary target *Mucilaginibacter* (arrows) obtained through combined pre- and probiotic treatment with both *Chitinophaga* and its niche prebiotic glutathione (Suppl. Figs. S12, S19); **d)** successful increase of primary targets *Burkholderia* and *Paenibacillus* and decrease of secondary targets *Rhizobium*, *Variovorax*, and *Bosea* (arrows) obtained through combined pre- and probiotic treatment with SynCom + ribose; **e)** successful increase of primary target *Burkholderia* and decrease of secondary targets *Rhizobium*, *Variovorax*, *Mucilaginibacter*, and *Chitinophaga* (arrows) obtained through prebiotic treatment (fructose); **d-e)** we also observed abundance changes of native soil members, in particular an increase of *Clostridium* (black triangle) when supplementing soil with sugars (see Suppl. Figs. S22-25); **f)** TE-based guild clustering measured in soil identifies *Streptomyces*, *Azospirillum*, and *Bosea* as *Clostridium*’s main competitors; **g-i)** fold-change of relative abundance (CPM) of *Clostridium* in soil against its main competitors *Azospirillum* (**a**), *Bosea* (**b**), and *Streptomyces* (**c**) across 30 pre- and/or probiotic conditions. Increasing *Clostridium*’s relative abundance in soil consistently decreased its main competitors (lower right quadrants) (see Suppl. Fig. S26).

We first sought to increase specific members (primary targets) of the SynCom present in soil and decrease their competitors (secondary targets), either through probiotic intervention (by adding either a single strain or the SynCom consortium), prebiotic intervention (supplementing the soil with primary target’s niche prebiotic, Suppl. Fig. S19), or combinations of both pre- and probiotic interventions. Our results confirmed that such MiND and guild predictions efficiently allowed us to design tailored pre- and probiotic interventions to induce targeted changes. For example, we selected to increase *Burkholderia* (primary target) accompanied by a decrease of its competitors *Rhizobium*, *Variovorax*, *Mucilaginibacter*, and *Chitinophaga* (secondary targets) (Suppl. Fig. S20). Our results demonstrated that combined pre- and probiotic treatments as well as prebiotic-only treatments significantly increased *Burkholderia* (up to 23-fold) and decreased its competitors (Fig. 4d,e, Suppl. Figs. S20, S22a-c, S23a-c, S24a-d, S25a-f), while probiotic-only treatment did not result in an increase of *Burkholderia* (Suppl. Fig. S21a). Similarly, we successfully increased *Paenibacillus* through prebiotics and combined pre- and probiotics treatments, but not by probiotic-only intervention (Fig. 4d, Suppl. Figs. S23e,f, S24b, S25b). Next, we targeted an increase of *Chitinophaga* and a decrease of its closest competitor *Mucilaginibacter*. Both probiotic (*Chitinophaga*) and combined pre- and probiotic (*Chitinophaga* + glutathione) treatments significantly increased *Chitinophaga* (fold- change = 7 and 13, respectively), while significantly decreasing its main competitor *Mucilaginibacter* (Fig. 4c, Suppl. Figs. S21c, S22d,e, S23d). A prebiotic-only treatment did not increase *Chitinophaga* in soil, probably because of *Chitinophaga’*s very low initial abundance (<50 CPM, Suppl. Fig. S24e,g).

Overall, we successfully increased primary targets in 4/7 probiotic-only treatments (Suppl. Fig S21), 12/14 combined pre- and probiotic treatments (using either a single strain or the SynCom consortium, Suppl. Figs. S22-24), and 7/7 prebiotic-only treatments (Suppl. Fig. S25). In over 95% (22/23) of cases in which a primary target increased, at least one of its competitors (secondary targets) decreased (Table 1, Suppl. Fig. S24). Of note, combined pre- and probiotic, or prebiotic-ony treatments often outperformed results from probiotic-alone treatments (Suppl. Fig. S23).

**Table 1.** Summary of targeted interventions carried out in soil.

| Intervention in soil | Total number of tested conditions | Number of conditions in which primary targets are increased | Number of conditions in which secondary targets are decreased |
| --- | --- | --- | --- |
| Probiotic (Single strain) | 7 | 4/7 (57%) | 4/4 (100%) |
| Prebiotic | 7 | 6/7 (86%) | 6/6 (100%) |
| Prebiotic + Probiotic (Single strain) | 10 | 8/10 (80%) | 6/8 (75%) |
| Prebiotic + Probiotic (Consortium) | 7 | 7/7 (100 %) | 7/7 (100%) |

Lastly, we explored if guild association can explain responses of non-SynCom microorganisms in soil to various pre- and probiotic interventions. We observed a strong increase in relative abundance of *Clostridium* accompanied by a decrease of *Azospirillum* upon addition of maltose + maltodextrin, trehalose, fructose, or ribose (Suppl. Fig. S25, Fig. 4d,e, Suppl. Figs. S22a- c,f,g, S23a-d, S24a-d). We calculated the TE for all microbes to predict competitive interactions in native soil and identified *Azospirillum*, *Bosea*, and *Streptomyces* as *Clostridium*’s main competitors (Fig. 4f). We observed a strong negative correlation between *Clostridium* and its main competitors across all 30 experiments, confirming that competition scores computed from guild associations explain responses of non-SynCom microbes to interventions in the community (Fig. 4g, Suppl. Fig. S26). Overall, TE-based guild classification and MiND can effectively predict competitive interactions of microorganisms in complex environments and can aid in the rapid design of interventions that selectively manipulate the microbiome.

### Broad applicability of TE-based guild classification

To demonstrate the broad applicability of this approach beyond the rhizosphere and soil, we measured TE and performed bacterial guild classification in fecal samples of healthy humans (n = 7, Suppl. Fig. S28). Our data revealed for example that *Faecalibacterium prausnitzii* and *Bifidobacterium longum* share the same guild and are competitors (positive competition scores) in all samples *B. longum* was detected (5/7). Although not significant, likely due to the small sample size, we observed that *F. prausnitzii* and *B. longum*’s relative abundance is negatively correlated (Spearman’s correlation coefficient *r* = -0.34). This corroborates results obtained by a study of 344 children and their response to a microbiota-directed complementary food intervention in which *F. prausnitzii* increased, while *B. longum* decreased^29, 30^.

## Discussion

Here, we present a novel approach that integrates *in vivo* measurements of transcription and translation to determine TE as a direct readout for each microbe’s prioritization for resource allocation^18^. The results are accurate predictions of competitive interactions and determination of substrate preferences, thus enabling effective intervention designs to selectively change complex microbiomes. While culture-independent approaches for microbiome studies often require hundreds to thousands of measurements to define correlation-based outcomes, our method generates a comprehensive understanding of microbial interactions and causality-based intervention strategies with just a single or a few experiments. Furthermore, our method provides an advantage over culture-dependent approaches that require isolates and are often limited by the number of combinations (e.g. pairwise) to be tested in diverse communities^12^. It is noteworthy that measurements taken in community settings, such as substrate utilization and TE, differed substantially from axenic measurements. For example, bacteria have the ability to utilize a number of substrates in axenic cultures but only prioritize a fraction of those when growing in a community. This suggests that determination of resource allocation based on axenic, culture-dependent experiments cannot always predict and explain the organism’s functionality when present in complex communities.

Our guild and niche determination predicted changes of the community to perturbation with high accuracy. The majority of those changes (81%) were attributed to substrate preferences (i.e. niche) or competition (i.e. guild) between community members (Suppl. Fig. S15). However, positive interactions between members of different guilds currently not accounted for by this approach might help to explain part of the remaining 19% of interactions^31^.

We revealed that guild associations are overall comparable between the SynCom and the SynCom in soil, explaining the high success rate of interventions in soil (Table 1). To address if these associations are also stable over time, we evaluated guilds in the SynCom after 4 and 7 days of incubation. We found that organisms are associated with similar guilds over time and that guild-based competition scores after 4 and 7 days were highly correlated (Pearson’s correlation coefficient *r* = 0.77, p-value < 2.2x10^-16^) (Suppl. Fig. S29).

MiND and guild analyses predict which SynCom members are likely to increase in relative abundance upon substrate addition and how competition influences the outcome of interventions. However, if two competitors in the same guild both prioritize the import of a specific substrate, e.g. *Burkholderia* and *Rhizobium* both prioritize ribose import (Fig. 3a), MiND could not predict the winner of this competition. To further assess competition outcome on a given substrate, we deployed genome-scale metabolic models (GEMs) of the 16 SynCom members to simulate growth with and without addition of selected substrates (see methods, Suppl. Table S4). GEMs predicted that both *Burkholderia* and *Rhizobium* would have a higher growth rate upon addition of ribose, which was confirmed by Biolog phenotypic microarray and growth curves in axenic cultures (Suppl. Table S3, S4, Suppl. Figs. S13, S14). However, GEMs also predicted that *Burkholderia* would outcompete *Rhizobium* because of *Burkholderia*’s higher growth rate upon ribose addition (Supp. Table S4). Furthermore, the models accurately simulated the effect of ribose addition on the other primary targets (high TE for ribose import proteins) *Paenibacillus* (increased), *Arthobacter* and *Brevibacillus* (no change) (Fig. 3e, Suppl. Table S4). While growth rate measurements can help to predict the winner in a competition of isolates, GEMs could assist to increase specificity and accuracy of MiND predictions for uncultivated microorganisms.

However, it is important to note that both GEMs as well as MiND-based design of prebiotic interventions rely on information about importer proteins. Availability of high quality annotations of transporters and their specificity, especially for environmental bacteria, is currently sparse^32, 33^. Therefore, well-curated genome annotations, as available for human gut microorganisms^34, 35^, will benefit the targeted design of prebiotic interventions.

Overall, guild elucidation and MiND explain a large percentage of community interactions and thus provide new insights into the functioning of biological systems. This understanding will be crucial for our ability to control and design microbiomes. Our approach will also help to identify targets for microbiome engineering, e.g. by CRISPR-Cas^36^, and will ultimately open the door to selectively alter microbial communities in a variety of environments, from aquatic and terrestrial to host-associated, and for a range of different applications, including environmental and human health related^37, 38^.

## METHODS

### Isolates

We established a 16-member microbial SynCom from the rhizosphere of switchgrass (*Panicum virgatum*) from agricultural crops, consisting of one strain each of *Arthrobacter*, *Bosea*, *Bradyrhizobium*, *Brevibacillus*, *Burkholderia*, *Chitinophaga*, *Lysobacter*, *Marmoricola*, *Methylobacterium*, *Mucilaginibacter*, *Mycobacterium*, *Niastella*, *Paenibacillus*, *Rhizobium*, *Rhodococcus* and *Variovorax*^19^. These isolates were obtained from the rhizosphere and soil surrounding a single switchgrass plant grown in marginal soils described elsewhere^39, 40^. Isolates and details on their isolation are available from the Leibniz Institute German Collection of Microorganisms and Cell Cultures GmbH (DSMZ) under accession numbers DSM 113524 (*Arthrobacter* OAP107), DSM 113628 (Bosea OAE506), DSM 113701 (*Bradyrhizobium* OAE829), DSM 113525 (*Brevibacillus* OAP136), DSM 113627 (Burkholderia OAS925), DSM 113563 (*Chitinophaga* OAE865), DSM 113522 (*Lysobacter* OAE881), DSM 114042 (*Marmoricola* OAE513), DSM 113562 (*Mucilaginibacter* OAE612), DSM 113602 (*Methylobacterium* OAE515), DSM 113539 (*Mycobacterium* OAE908), DSM 113593 (*Niastella* OAS944), DSM 113526 (*Paenibacillus* OAE614), DSM 113517 (*Rhizobium* OAE497), DSM 113518 (*Rhodococcus* OAS809), DSM 113622 (*Variovorax* OAS795).

### Isolates growth conditions

Precultures of individual isolates were generated in 5 mL of liquid 1x R2A medium (Teknova, cat # R0005) under oxic conditions and grown at 30 °C for 7 days without shaking. One isolate (*Bradyrhizobium* OAE829) was grown in 0.1x R2A due to poor growth in 1x R2A, as previously described^19^.

### SynCom assembly and growth conditions

Optical density readings at 600 nm (OD600), from pre-cultures were taken by a Molecular Devices SpectraMax M3 Multi-Mode Microplate Reader (VWR, cat # 89429-536). Pre-cultures were diluted to a starting OD600 of 0.02 in 5 mL 0.1x R2A. The SynCom was assembled in large volumes to minimize pipetting error and maximize reproducibility^19^. Briefly, 1 mL of each normalized culture (OD600 = 0.02) was diluted in a final volume of 250 mL 0.1x R2A and spread into 20 mL aliquots in Falcon tubes. Falcon tubes containing the SynCom inoculum were then incubated at 30 °C for 7 days aerobically in four biological replicates. After 7 days of growth the SynCom samples were harvested by centrifugation and pellets were immediately treated for multi-omics analysis as detailed below (DNA-, RNA- and metaRibo-Seq).

### Targeted interventions in the SynCom

For targeted modification experiments in the SynCom, the SynCom was assembled and grown as described above. Specific isolates were omitted from the SynCom assembly for the dropout experiments, as described in Fig. 2e. Alternatively, concentrated stocks of either *Burkholderia*, *Chitinophaga*, *Mucilaginibacter*, or *Rhizobium* were added as probiotics to the SynCom, as described in Suppl. Fig. S7.

Metabolites from concentrated, filter-sterilized stocks of compounds were added to the medium for the prebiotics experiments, as described in Fig. 3e-j, Suppl. Figs. S15 and S16. Initially, a total of 14 compounds were tested in three different concentrations, including six sugars (fructose, galactose, maltose/maltodextrin, ribose, trehalose, xylose), three amino acids (cystine, glutamate, methionine), two diamines (putrescine, spermidine), one vitamin (cobalamin), one peptide (glutathione), and one inorganic compound (sulfate/thiosulfate) (Suppl. Fig. S15-S16). Addition of cobalamin, cystine, or methionine did not induce significant change in relative abundance in the community, likely because these substrates were already present in excess in the non-modified culture medium; we thus discarded these three compounds from subsequent analysis. Experiments were carried out in duplicates and pellets were stored at -80 °C prior to metagenomic analysis.

### Natural soil incubation and growth conditions

Root associated soil was collected from the Oklahoma State University research farm near Perkins, OK, USA (35.991148, -97.046489, elevation 280 m). The soil surface was cleaned of all organic debris. Soil and roots from *Panicum virgatum* (i.e. switchgrass) were obtained by shovel to a depth of 20 cm immediately adjacent to the crown margin of the target switchgrass plant. Samples were quickly frozen after sampling and stored at -20 °C prior to use.

Fifty gram (50 g) of frozen soil was added to 250 mL of 0.1x R2A culture medium or 0.1x R2A + SynCom inoculum prepared as described above, and this volume was distributed (5 mL each) into 14 mL culture tubes. We then added 50-500 µL of concentrated, filter-sterilized stocks of compounds (similar to a prebiotic treatment) and/or 20 µL of the SynCom isolates diluted at an OD600 of 0.02 (i.e. approximately 8x10^5^ CFU/mL) (similar to a probiotic treatment) as described in Fig. 4a. Soil samples were grown at 30 °C for 7 days aerobically in duplicates (three replicates were used for the soil reference sample), harvested by centrifugation and pellets were stored at -80 °C prior to metagenomic analysis.

### Human fecal sample collection and processing

Volunteers were recruited in accordance with the institutional review board (IRB) number 150275. Inclusion criteria were: no known medical condition or treatment during the past three months, and no antibiotic treatment over the past six months. Fecal samples from seven self-described healthy individuals were collected and immediately frozen at -80 °C (<2 min after collection).

### Metagenomic (DNA-Seq) and metatranscriptomic (RNA-Seq) sample preparation

DNA and RNA from SynCom samples were extracted from leftover lysates from metaRibo-Seq sample preparation (see below) and stored in Trizol at -80 °C. DNA from soil samples was extracted using the ZymoBIOMICS DNA miniprep kit (Zymo). RNA was extracted using a RNeasy mini kit (Qiagen) and rRNA was removed using QIAseq FastSelect-5S/16S/23S kit (Qiagen). DNA and RNA from fecal samples were extracted using the ZymoBIOMICS DNA/RNA miniprep kit (Zymo). DNA-Seq libraries were prepared using Nextera XT library preparation kit with 700 pg DNA input per sample and 6:30 min tagmentation at 55 °C and barcoded using Nextera XT indexes (Illumina). RNA-Seq libraries were prepared using KAPA RNA HyperPrep kit (Roche) and barcoded using TruSeq indexes (Illumina). Amplification was followed in real time using SYBR-Green and stopped when reaching a plateau.

### Metatranslatomic (metaRibo-Seq) sample preparation

Metatranslatomic (metaRibo-Seq) sample preparations were performed according to the protocol provided in Suppl. Material 1. This protocol is based on a previously published Ribo-Seq protocol for axenic bacterial cultures^23^ and shares similarities with a recently published MetaRibo-Seq protocol from Fremin et al^41^. Briefly, mechanical bacterial lysis was performed in a solution containing chloramphenicol and Guanosine-5’-[(β,γ)-imido]triphosphate (GMPPNP) to stop protein elongation. Resulting lysates were treated with MNase and DNase to degrade nucleic acids that were not protected by ribosomes. Monosome recovery was performed using RNeasy Mini spin size-exclusion columns (Qiagen) and RNA Clean & Concentrator-5 kit (Zymo). rRNA removal was performed using the QIAseq FastSelect- 5S/16S/23S kit (Qiagen). MetaRibo-Seq libraries were prepared using the NEBNext Small RNA Library Prep set for Illumina, with modifications (see details in Suppl. Material 1). Amplification was followed in real time using SYBR-Green and stopped when reaching a plateau. PCR products were purified using Select-a-size DNA Clear & Concentrator kit (Zymo). Leftover lysate prior to MNase treatment was saved and stored at -80 °C for metagenomic and metatranscriptomic analysis.

### Sequencing

The quality and average size of the libraries was controlled using a 4200 TapeStation System (Agilent). Library concentrations were quantified using Qubit dsDNA HS Assay kit and QuBit 2.0 Fluorometer (Invitrogen). Libraries were sequenced on an Illumina NovaSeq, PE100 platform. Minimum sequencing depth was 10 million reads for metagenomic samples, 50 million reads for metatranscriptomic samples, and 100 million reads for metatranslatomic samples.

### Reference genomes for SynCom members

Genomic data from individual cultures of the 16 SynCom members was used to assemble genomes using SPAdes version /3.13.0^42^ and quality controlled using CheckM version v1.0.13^43^. 16S rRNA phylogenetic tree analysis was performed using Clustal Omega^44^. Genomes were annotated at the gene level using PROKKA version 1.14.5^45^ and KEGG pathway annotation was performed using BlastKOALA version 2.2^46^. A custom SynCom metagenome database was built from the genomes of the 16 isolates using bowtie2 version 2.3.2^47^.

### Databases

Data from soil microbiome samples were aligned to a modified Web of Life (WoL) database^35^. Modification of the WoL database was performed to reduce false alignment hits. Briefly, we calculated genome coverages in the metagenomic samples using Zebra^48^. We then created a reduced version of the WoL database including only genomes with at least 50% aggregated coverage across all soil experiments. Multi-omic data (i.e. metagenomic, metatranscriptomic, metaRibo-Seq) from the soil experiments were aligned to the modified WoL database. Human-associated microbial genomes are generally better referenced in public databases, thus data from human gut microbiome samples were aligned to WoL directly.

### Data processing

Adapter sequences were removed from multi-omics sequencing data using TrimGalore (Cutadapt) version 1.18^49^ and quality controlled using FastQC version 0.11.9^50^. Trimmed reads were aligned to the appropriate database using bowtie2 version 2.3.2^47^. Gene and KEGG pathway count tables stratified by genus were obtained using Woltka version 0.1.1^35^. Multi- omics gene counts were normalized to reads per kilobase per million (RPKM) for the SynCom experiments, or counts per million (CPM) for soil and gut microbiome samples.

### Statistics

Feature count tables were imported and analyzed using R version 3.6.3^51^. Hierarchical clustering on the principal components (HCPC) analysis was performed using the FactomineR package^52^.

### TE calculation

We calculated TE as the ratio between Ribo-Seq and RNA-Seq signal as follows:

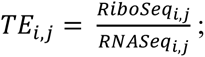

Where *RiboSeq_i,j_* and *RNASeq_i,j_* are Ribo-Seq and RNA-Seq normalized read counts for each *feature i* in each *bacteria j*.

### Metabolic guild clustering

Microbial community members were classified into functional guilds based on TE measured on KEGG-annotated metabolic pathways by performing a hierarchical clustering on the principal components (HCPC) analysis. Briefly, the HCPC algorithm comprises 3 steps: *i)* principal component analysis (PCA), *ii)* hierarchical clustering on the principal components, and *iii)* k-mers partitioning to stabilize initial classification^52^.

### Competition scoring

For each pair of *bacteriai,j* in a microbial community we defined a *Competition scorei,j*, referring to the likeliness of a competition interaction between bacteria *i* and *j*, as a function of the distance between *i* and *j* in the TE-based guild clustering as follows:

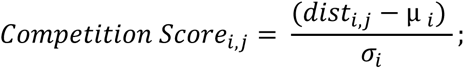

Where *disti,j* is the euclidean distance between bacteria *i* and *j* in the guild clustering, *μi* and *σi* are the average and the standard deviation of the population distances to bacteria *i.* Note: for predictions made in the SynCom, *Scorei,j* was adjusted to zero for bacteria having a relative abundance <0.5% after 7 days of growth.

Sensitivity and specificity of the prediction of community outcomes upon modifications of the community composition were calculated by considering a *Scorei,j* > 0 as “likely” and *Scorei,j* ≤ 0 as “unlikely” for bacteria *j* to increase/decrease upon removal/increase of bacteria *i* relative abundance in the community.

### Microbial Niche Determination (MiND)

Microbial niches were identified as metabolites for which a translational activity (positive TE) was measured in one or more microbial community members on import protein(s) for a given substrate.

### Definition of primary and secondary targets

Primary targets were defined as microbes which we sought to enrich in the community, either by probiotic or prebiotic addition with substrates identified as preferential to the microbe by MiND. Secondary targets were defined as microbes we sought to decrease in the community by promoting one or more of their competitors as defined based on guild clustering (their competitor(s) would then be considered as primary target).

### Phenotypic microarray assays for axenic cultures

Isolates for all 16 SynCom members were grown axenically on 285 different substrates using Biolog Phenotypic Microarray (PM) plates (PM 1, 2A, 3B) following the company’s instructions (Biolog). Briefly, isolates were streaked on 1x R2A agar plates (1.5% w/v) (*Bradyrhizobium* was streaked on 0.1x R2A plates), colonies were picked and resuspended in inoculation fluid IF-0a GN/GP (cat no. 72268) up to an OD600 of 0.07 and inoculated into the PM plates in triplicate. Plates were incubated at 30 °C without shaking with lids coated with an aqueous solution of 20% ethanol and 0.01% Triton X-100 (Sigma) to prevent condensation^19^. Growth in PM plates was indicated by color change from clear to purple of Biolog Redox Dye Mixes (Dye Mix G and H, cat nos. 74227, 74228). Additionally, we defined substrate utilization as OD600 increase >0.02 over a 7 day incubation period.

### Genome-scale metabolic models simulations

Genome-scale metabolic models (GEMs) of the 16 SynCom members were reconstructed. Models were simulated using Flux Balance Analysis (FBA)^53^. The predicted growth rates of SynCom members were recorded for comparison purposes. To predict the effect of media supplementation with different substrates, we incorporated the uptake flux of each substrate at a time and simulated the model for each condition (i.e. change in media composition). A list of the substrates provided in Suppl. Table S4 that contains the predicted growth rates tested in this analysis. All GEMs were analyzed using COBRApy software package version 0.17.1^54^ with IBM CPLEX solver version 22.1.0 (IBM) in Python (version 3.7.11).

## Supporting information

MetaRibosome Profiling Protocol

## Figures

Fig. 1a and 4a were created using BioRender.com. Other graphical representations were produced using R version 3.6.3^51^ and associated packages ggplot2, gplots version 3.0.1.1, cowplot version 1.1.1 and factoextra version 1.0.7 packages^55–58^.

## Data availability

Multi-omics sequencing data generated through this study are available on the NCBI Sequence Read Archive (submission SUB12797845, BioProject PRJNA942264).

## Code availability

R code used to perform statistical analysis presented in this manuscript is available at https://github.com/ZenglerLab/MIND-.

## Authors contributions

O.M., M.A.B. and K.Z. designed the study. O.M. supervised experiments, carried out bioinformatics data processing with help from M.A.B. and L.Z.; O.M. performed statistical analysis and generated figures. M.A.B. adapted the bacterial Ribo-Seq protocol published previously by our group to be extended to microbial communities with help from O.M.; O.M. performed cultivation experiments with help from C.L. All multi-omic experiments were carried out by O.M., M.A.B. and C.L. Spot-on-lawn competition assays were performed by G.J.N. and phenotypic microarray assays were carried out by D.T. Growth curve assay data and pathway annotation for all SynCom genomes were done by E.H. Genome-scale metabolic models of the SynCom members were reconstructed and curated by M.K. O.M. and K.Z. wrote the manuscript with input from all the co-authors.

## Funding

This material is based upon work supported by the U.S. Department of Energy, Office of Science, Office of Biological & Environmental Research under Awards DE-SC0021234 and DE-SC0022137. Furthermore, the development of the technologies described in this article were in part funded through Trial Ecosystem Advancement for Microbiome Science Program at Lawrence Berkeley National Laboratory funded by the U.S. Department of Energy, Office of Science, Office of Biological & Environmental Research Awards DE-AC02-05CH11231. The work was also supported by the UC San Diego Center for Microbiome Innovation (CMI) through a Grand Challenge Award. This publication includes data generated at the UC San Diego IGM Genomics Center utilizing an Illumina NovaSeq 6000 that was purchased with funding from a National Institutes of Health SIG grant (#S10 OD026929). G.J.N was supported in part by the NIH training grant T32 DK007202.

## Acknowledgments

We are thankful to Thomas Juenger and Jason Bonnette (University of Texas at Austin) for providing us with switchgrass soil samples for this study. We thank Daniel Hakim (UCSD) for assistance with Zebra filter analysis and Juan Tibocha-Bonilla (UCSD) for help with drafting figures. We are grateful to Kateryna Zhalnina (LBNL) and Adam Deutschbauer (LBNL) for assisting with genome assembly for the SynCom members. Furthermore, we would like to acknowledge fruitful discussions with Trent Northen (LBNL) and Rob Knight (UCSD) that helped improve an earlier version of the manuscript.

## Competing interest

O.M., M.A.B., and K.Z. are inventors on a related patent application.

**Supplementary Information is available for this paper.**

**Correspondence and requests for materials should be addressed to KZ (**).

## References

1. Leshem, A., Segal, E. & Elinav, E. The gut microbiome and individual-specific responses to diet. mSystems 5, (2020).

2. Zhalnina, K. et al. Dynamic root exudate chemistry and microbial substrate preferences drive patterns in rhizosphere microbial community assembly. Nat Microbiol 3, 470–480 (2018).

3. Zmora, N., Zeevi, D., Korem, T., Segal, E. & Elinav, E. Taking it personally: personalized utilization of the human microbiome in health and disease. Cell Host Microbe 19, 12–20 (2016).

4. Paoli, L. et al. Biosynthetic potential of the global ocean microbiome. Nature 607, 111– 118 (2022).

5. Bahram, M. et al. Structure and function of the global topsoil microbiome. Nature 560, 233–237 (2018).

6. Prosser, J. I. How and why in microbial ecology: An appeal for scientific aims, questions, hypotheses and theories. Environ. Microbiol. 24, 4973–4980 (2022).

7. Walter, J., Armet, A. M., Finlay, B. B. & Shanahan, F. Establishing or exaggerating causality for the gut microbiome: lessons from human microbiota-associated rodents. Cell 180, 221–232 (2020).

8. Milligan-McClellan, K. C. et al. Deciphering the microbiome: integrating theory, new technologies, and inclusive science. mSystems 7, e0058322 (2022).

9. Gilbert, J. A. et al. Current understanding of the human microbiome. Nat. Med. 24, 392– 400 (2018).

10. Vorholt, J. A., Vogel, C., Carlström, C. I. & Müller, D. B. Establishing causality: opportunities of synthetic communities for plant microbiome research. Cell Host Microbe 22, 142–155 (2017).

11. Trivedi, P., Leach, J. E., Tringe, S. G., Sa, T. & Singh, B. K. Plant-microbiome interactions: from community assembly to plant health. Nat. Rev. Microbiol. 18, 607–621 (2020).

12. Kehe, J. et al. Positive interactions are common among culturable bacteria. Sci Adv 7, eabi7159 (2021).

13. Wang, Z. et al. Complementary resource preferences spontaneously emerge in diauxic microbial communities. Nat. Commun. 12, 6661 (2021).

14. Jo, C., Bernstein, D. B., Vaisman, N., Frydman, H. M. & Segrè, D. Construction and modeling of a coculture microplate for real-time measurement of microbial interactions. mSystems e0001721 (2023).

15. Morin, M. A., Morrison, A. J., Harms, M. J. & Dutton, R. J. Higher-order interactions shape microbial interactions as microbial community complexity increases. Sci. Rep. 12, 22640 (2022).

16. Buttgereit, F. & Brand, M. D. A hierarchy of ATP-consuming processes in mammalian cells. Biochem. J 312 **(Pt** **1**), 163–167 (1995).

17. Ingolia, N. T. Ribosome profiling: new views of translation, from single codons to genome scale. Nat. Rev. Genet. 15, 205–213 (2014).

18. Al-Bassam, M. M. et al. Optimization of carbon and energy utilization through differential translational efficiency. Nat. Commun. 9, 4474 (2018).

19. Coker, J., et al. A reproducible and tunable synthetic soil microbial community provides new insights into microbial ecology. mSystems 7, e0095122 (2022).

20. Le Scornet, A. & Redder, P. Post-transcriptional control of virulence gene expression in *Staphylococcus aureus*. Biochim. Biophys. Acta Gene Regul. Mech. 1862, 734–741 (2019).

21. Fris, M. E. & Murphy, E. R. Riboregulators: Fine-tuning virulence in *Shigella*. Front. Cell. Infect. Microbiol. 6, 2 (2016).

22. Romby, P., Vandenesch, F. & Wagner, E. G. H. The role of RNAs in the regulation of virulence-gene expression. Curr. Opin. Microbiol. 9, 229–236 (2006).

23. Latif, H. et al. A streamlined ribosome profiling protocol for the characterization of microorganisms. Biotechniques 58, 329–332 (2015).

24. Fremin, B. J., Sberro, H. & Bhatt, A. S. MetaRibo-Seq measures translation in microbiomes. Nat. Commun. 11, 3268 (2020).

25. Kanehisa, M. & Goto, S. KEGG: kyoto encyclopedia of genes and genomes. Nucleic Acids Res. 28, 27–30 (2000).

26. Root, R. B. The niche exploitation pattern of the blue-gray gnatcatcher. Ecol. Monogr. 37, 317–350 (1967).

27. Simberloff, D. & Dayan, T. The guild concept and the structure of ecological communities. Annu. Rev. Ecol. Syst. 22, 115–143 (1991).

28. Durand, G. A., Raoult, D. & Dubourg, G. Antibiotic discovery: history, methods and perspectives. Int. J. Antimicrob. Agents 53, 371–382 (2019).

29. Gehrig, J. L. et al. Effects of microbiota-directed foods in gnotobiotic animals and undernourished children. Science 365, eaau4732 (2019).

30. Raman, A. S. et al. A sparse covarying unit that describes healthy and impaired human gut microbiota development. Science 365, eaau4735 (2019).

31. Zuñiga, C. et al. Environmental stimuli drive a transition from cooperation to competition in synthetic phototrophic communities. Nat Microbiol 4, 2184–2191 (2019).

32. Saier, M. H., Jr et al. The Transporter Classification Database (TCDB): recent advances. Nucleic Acids Res. 44, D372–9 (2016).

33. Kermani, A. A. et al. The structural basis of promiscuity in small multidrug resistance transporters. Nat. Commun. 11, 6064 (2020).

34. Franzosa, E. A. et al. Species-level functional profiling of metagenomes and metatranscriptomes. Nat. Methods 15, 962–968 (2018).

35. Zhu, Q. et al. Phylogenomics of 10,575 genomes reveals evolutionary proximity between domains Bacteria and Archaea. Nat. Commun. 10, 5477 (2019).

36. Rubin, B. E. et al. Species- and site-specific genome editing in complex bacterial communities. Nat Microbiol 7, 34–47 (2022).

37. Sheth, R. U., Cabral, V., Chen, S. P. & Wang, H. H. Manipulating bacterial communities by *in situ* microbiome engineering. Trends Genet. 32, 189–200 (2016).

38. Zaramela, L. S. et al. The sum is greater than the parts: exploiting microbial communities to achieve complex functions. Curr. Opin. Biotechnol. 67, 149–157 (2021).

39. Sher, Y. et al. Microbial extracellular polysaccharide production and aggregate stability controlled by switchgrass (*Panicum virgatum*) root biomass and soil water potential. Soil Biol. Biochem. 143, 107742 (2020).

40. Ceja-Navarro, J. A. et al. Protist diversity and community complexity in the rhizosphere of switchgrass are dynamic as plants develop. Microbiome 9, 96 (2021).

41. Fremin, B. J., Nicolaou, C. & Bhatt, A. S. Simultaneous ribosome profiling of hundreds of microbes from the human microbiome. Nat. Protoc. 16, 4676–4691 (2021).

42. Bankevich, A. et al. SPAdes: a new genome assembly algorithm and its applications to single-cell sequencing. J. Comput. Biol. 19, 455–477 (2012).

43. Parks, D. H., Imelfort, M., Skennerton, C. T., Hugenholtz, P. & Tyson, G. W. CheckM: assessing the quality of microbial genomes recovered from isolates, single cells, and metagenomes. Genome Res. 25, 1043–1055 (2015).

44. Sievers, F. & Higgins, D. G. The clustal omega multiple alignment package. Methods Mol. Biol. 2231, 3–16 (2021).

45. Seemann, T. Prokka: rapid prokaryotic genome annotation. Bioinformatics 30, 2068–2069 (2014).

46. Kanehisa, M., Sato, Y. & Morishima, K. BlastKOALA and GhostKOALA: KEGG tools for functional characterization of genome and metagenome sequences. J. Mol. Biol. 428, 726–731 (2016).

47. Langmead, B. & Salzberg, S. L. Fast gapped-read alignment with Bowtie 2. Nat. Methods 9, 357–359 (2012).

48. Hakim, D. et al. Zebra: Static and dynamic genome cover thresholds with overlapping references. mSystems 7, e0075822 (2022).

49. Martin, M. Cutadapt removes adapter sequences from high-throughput sequencing reads. EMBnet.journal 17, 10–12 (2011).

50. Andrews, S. FastQC: a quality control tool for high throughput sequence data. Babraham bioinformatics version 0.11. 7. Preprint at (2010).

51. Core Team, R. R: A language and environment for statistical computing. Version 3.6. 0. Vienna, Austria. */ra-language-and-environment-forstatistical-computing*.

52. Lê, S., Josse, J. & Husson, F. FactoMineR: An R package for multivariate analysis. J. Stat. Softw. 25, 1–18 (2008).

53. Orth, J. D., Thiele, I. & Palsson, B. Ø. What is flux balance analysis? Nat. Biotechnol. 28, 245–248 (2010).

54. Ebrahim, A., Lerman, J. A., Palsson, B. O. & Hyduke, D. R. COBRApy: COnstraints- based reconstruction and analysis for python. BMC Syst. Biol. 7, 74 (2013).

55. Wickham, H. ggplot2: elegant graphics for data analysis New York. NY: Springer.

56. Warnes, G. R., Bolker, B., Bonebakker, L. & Gentleman, R. gplots: various R programming tools for plotting data, version 3.0. 1. Search in.

57. Wilke, C. O. cowplot: streamlined plot theme and plot annotations for ‘ggplot2’ https://CRAN.R-project.org/package=cowplot.

58. Kassambara, A. & Mundt, F. Factoextra: extract and visualize the results of multivariate data analyses, R package version 1.0. 7. 2020. Preprint at (2021).

