## Supplementary Figures and Tables for "Guild and Niche Determination Enable Targeted Alteration of the Microbiome"

### EXTENDED DATA FIGURES

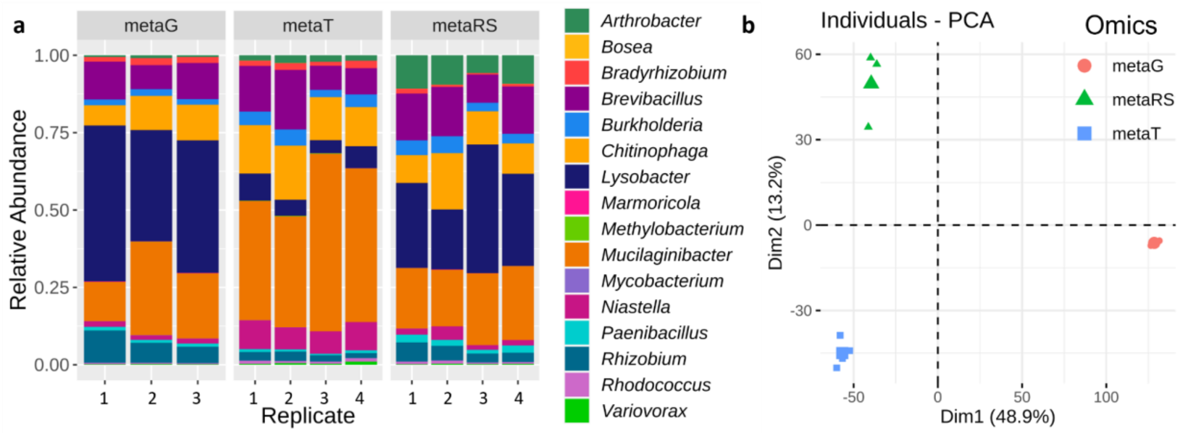


**Suppl. Fig. S1. Reproducibility of multi-omics analysis across biological replicates. a)** Relative abundance of the 16 SynCom members across replicates at the metagenomic (metaG), metatranscriptomic (metaT), and metatranslatomic (metaRS) level; **b)** PCA of the metagenomic, metatranscriptomic, and metatranslatomic profiles on >11,000 Strain|gene features (log-transformed RPKM). Bigger symbols represent the average for each group.

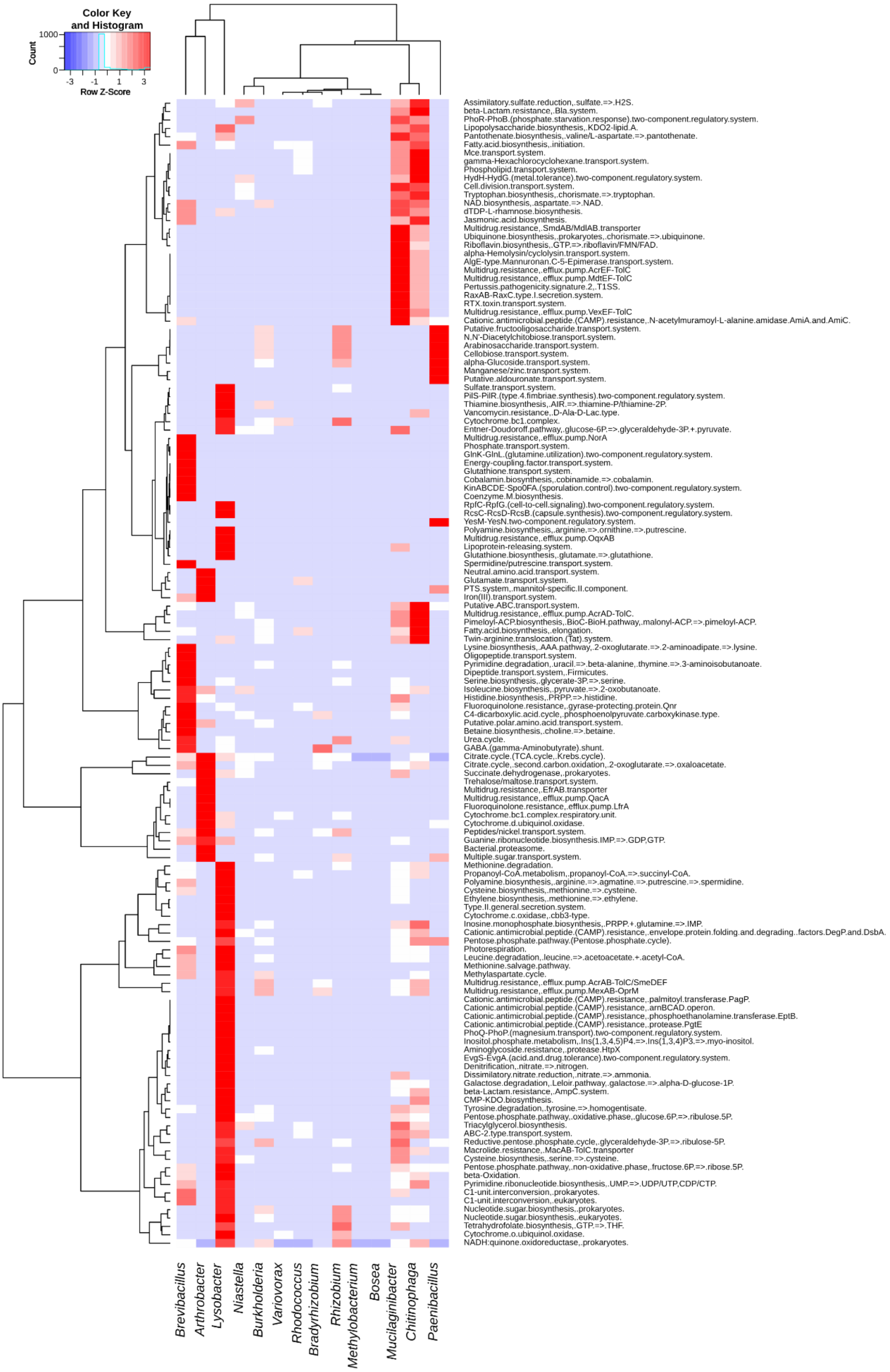


**Suppl. Fig. S2. Differential metabolic pathway prioritization in the SynCom guilds.** KEGG pathways that are significantly associated (see methods) to at least one guild are shown. Guild clustering as defined in Fig. 1b,c is shown on top. See also Suppl. Table S1 for numeric information.

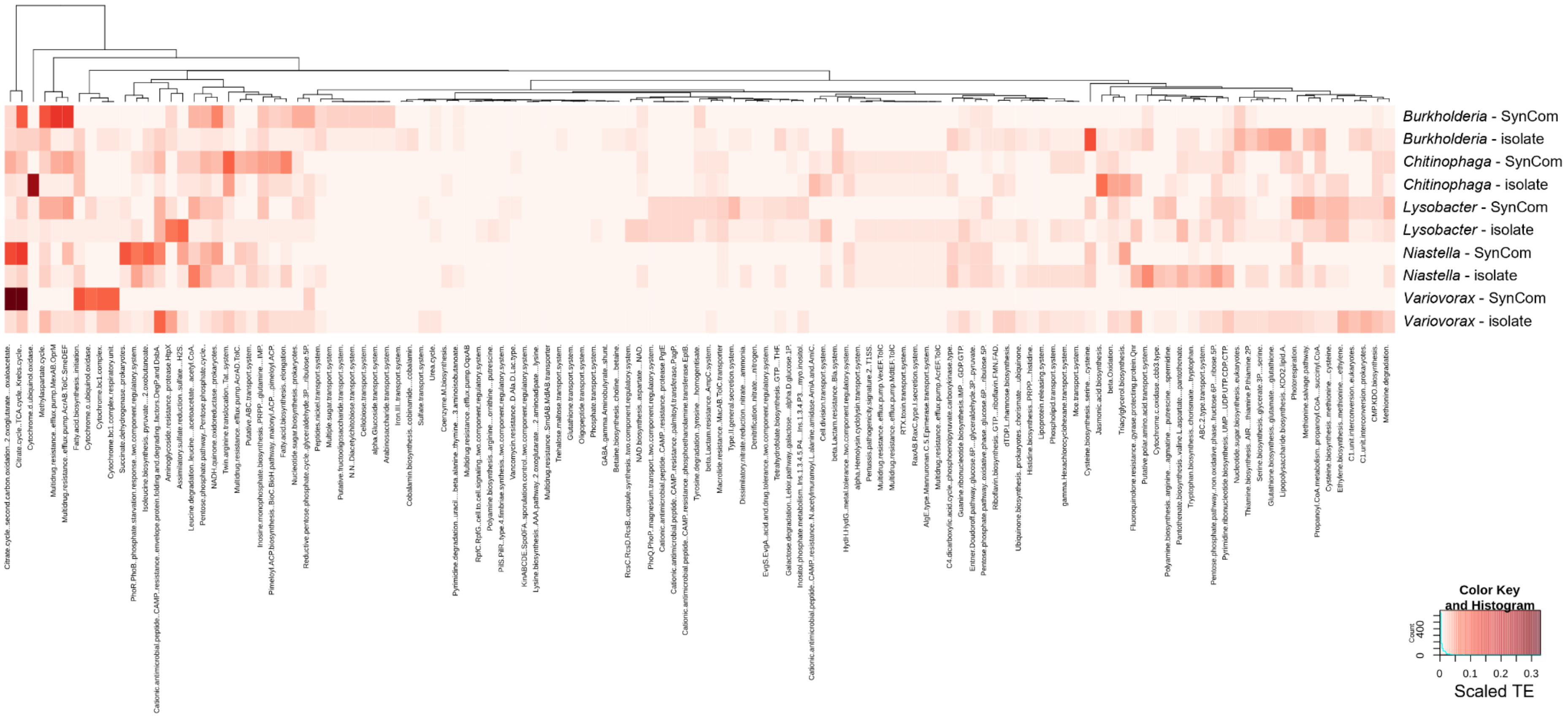


**Suppl. Fig. S3. Metabolic pathway prioritization in the microbial community is different than in axenic culture.** Heatmap of TE (row-scaled) in 137 KEGG pathway that significantly discriminate the metabolic guilds as defined in Fig. 1 (see methods, Fig. 1) for microbes grown in the SynCom (n = 4) or in axenic cultures (n = 3) in the same condition (7 days at 30 °C in 0.1x R2A).

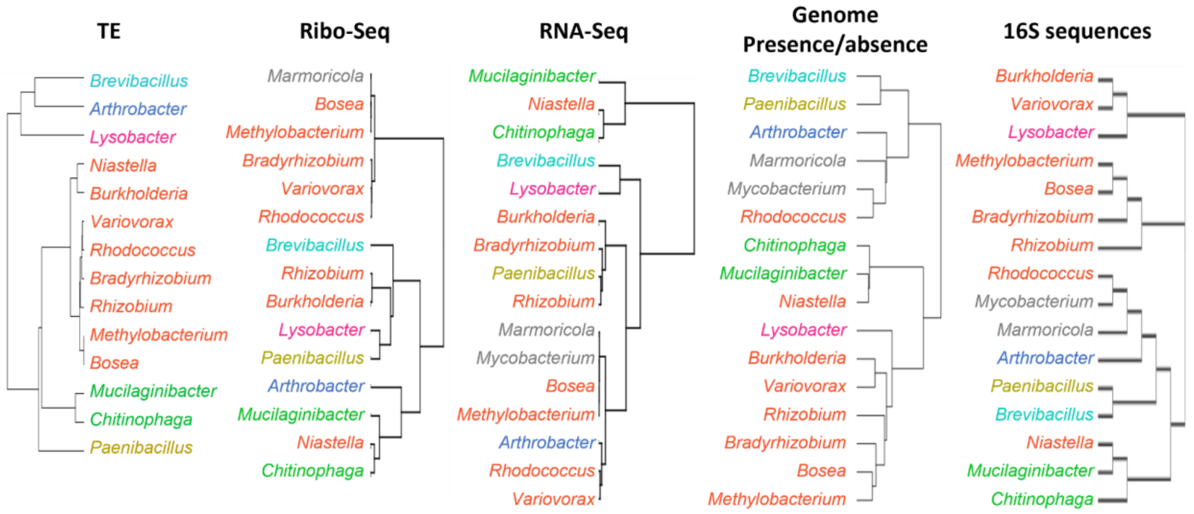


**Suppl. Fig. S4. Guild-based microbiome classification based on metabolic pathways prioritization (TE) differ from classifications based on Ribo-Seq, RNA-Seq, presence/absence of pathways in the genome, or 16S rRNA-based phylogeny.** All analyses were based on RPKM counts aggregated by KEGG pathways. Color coding of metabolic guilds as defined in Fig. 1b,c has been kept for comparison.

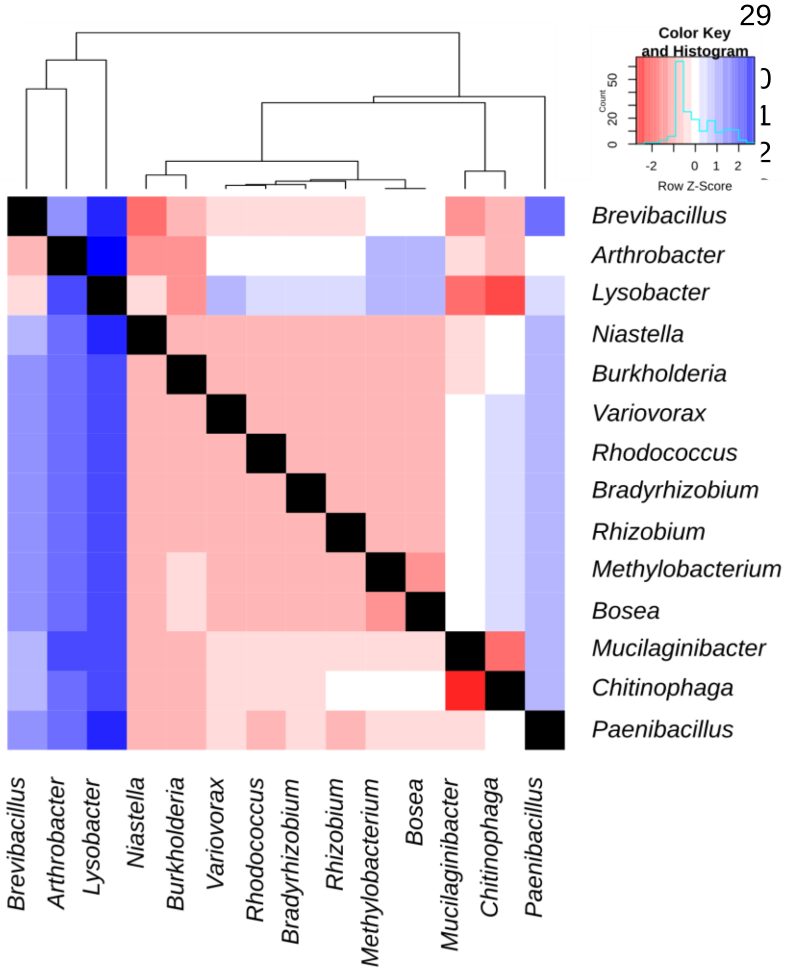


**Suppl. Fig. S5. Dendrogram and distance matrix of TE-based guild clustering (row Z-score).** Values were used as a basis to build the competition score shown in Fig. 2d.

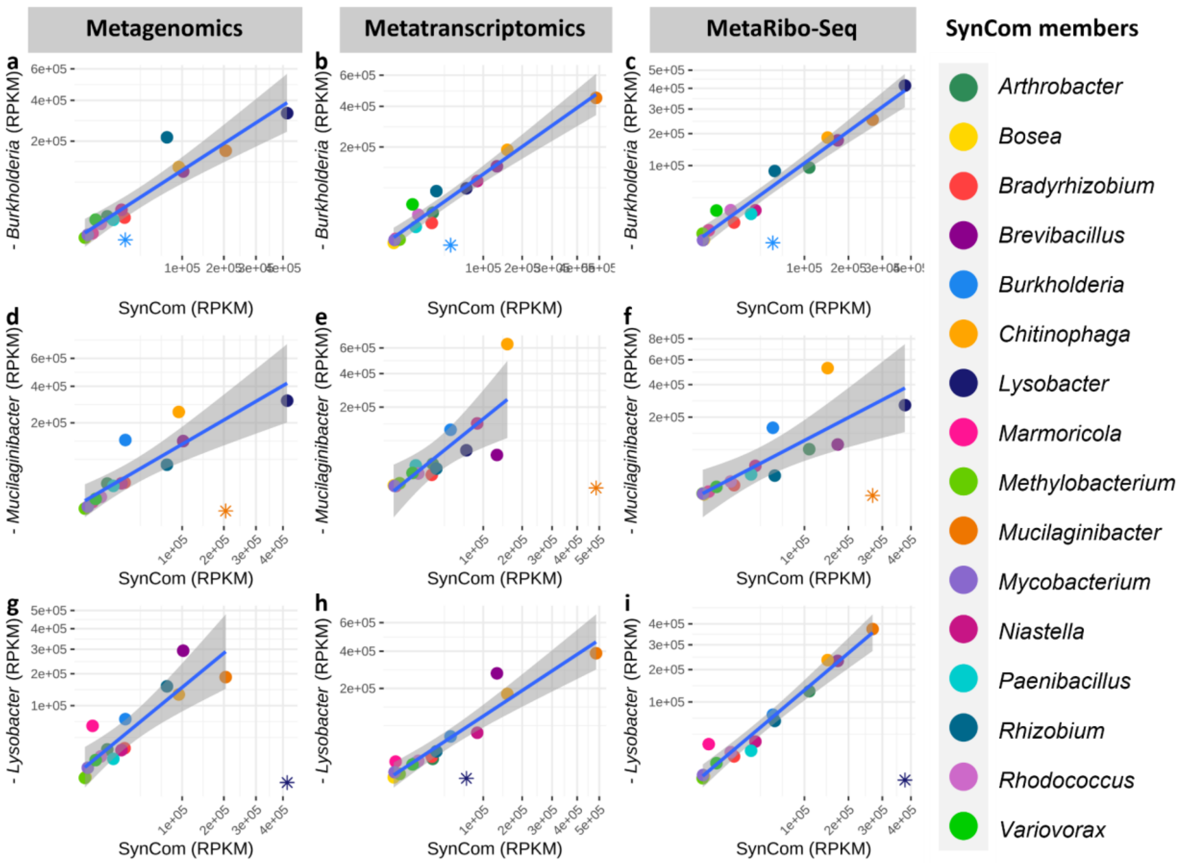


**Suppl. Fig. S6. Single-strain removal increases relative abundance, transcriptional and translational activity of guild-predicted competitors in the SynCom.** Multi-omics taxonomy analysis of single-strain dropout experiments in the SynCom. Linear regression and 99% confidence interval (CI) of square-transformed RPKM in SynCom control (x axis) vs. dropout SynCom (y axis). Organisms above the 99% CI were considered significantly increased upon removal of the dropped out member; Top to bottom: *Burkholderia* (**a-c**), *Mucilaginibacter* (**d-f**), and *Lysobacter* (**g-i**) dropout experiments. Left to right: metagenomic (**a,d,g**), metatranscriptomic (**b,e,h**) and metatranslatomic (**c, f, i**) taxonomy analysis.

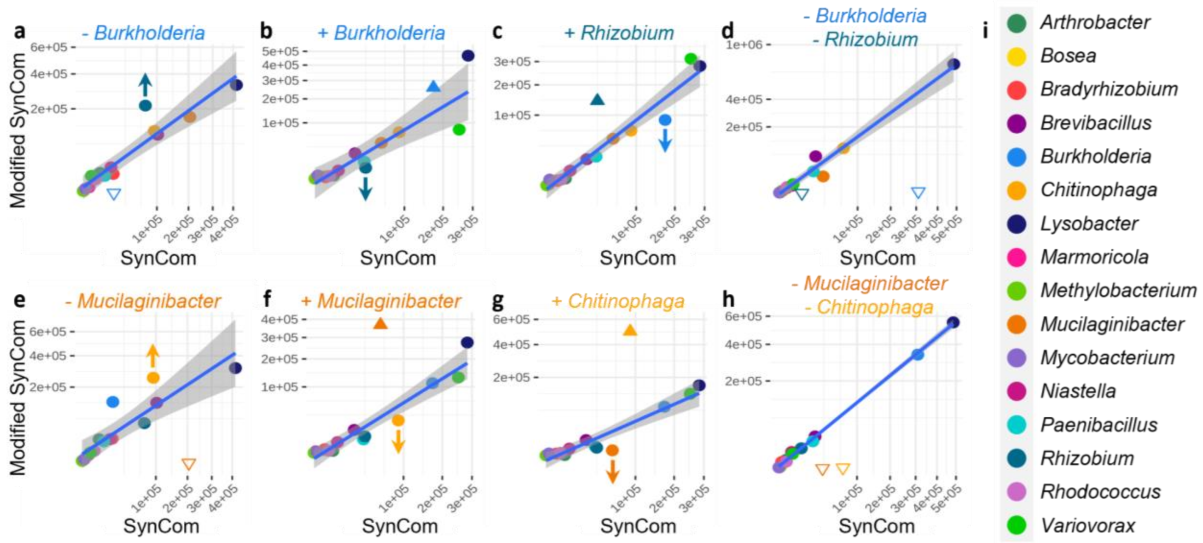


**Suppl. Fig. S7. In-detail competition assays of two close competition pairs**. *Burkholderia-Rhizobium* and *Mucilaginibacter-Chitinophaga* were predicted to be strong guild competitors in the SynCom (Fig. 2d); **a-h)** linear regression and 99% confidence interval (CI) of square-transformed RPKM in control (x axis) vs. modified community (y axis). Organisms above/below the 99% CI were considered significantly increased/decreased in the modified community; **a-d)** *Burkholderia-Rhizobium* competition testings: **a)** *Burkholderia* dropout increased *Rhizobium;* **b)** *Burkholderia* (20x) enrichment decreased *Rhizobium*; **c)** *Rhizobium* (20x) enrichment decreased *Burkholderia*; **d)** *Burkholderia-Rhizobium* double dropout had very minor effect on the composition of the rest of the community; **e-h)** *Burkholderia-Rhizobium* competition testings: **e)** *Mucilaginibacter* dropout increased *Chitinophaga;* **f)** *Mucilaginibacter* (100x) enrichment decreased *Chitinophaga*; **g)** *Chitinophaga* (100x) enrichment decreased *Mucilaginibacter*; **h)** *Mucilaginibacter-Chitinophaga* double dropout did not have any effect on the composition of the rest of the community; **i)** color key.

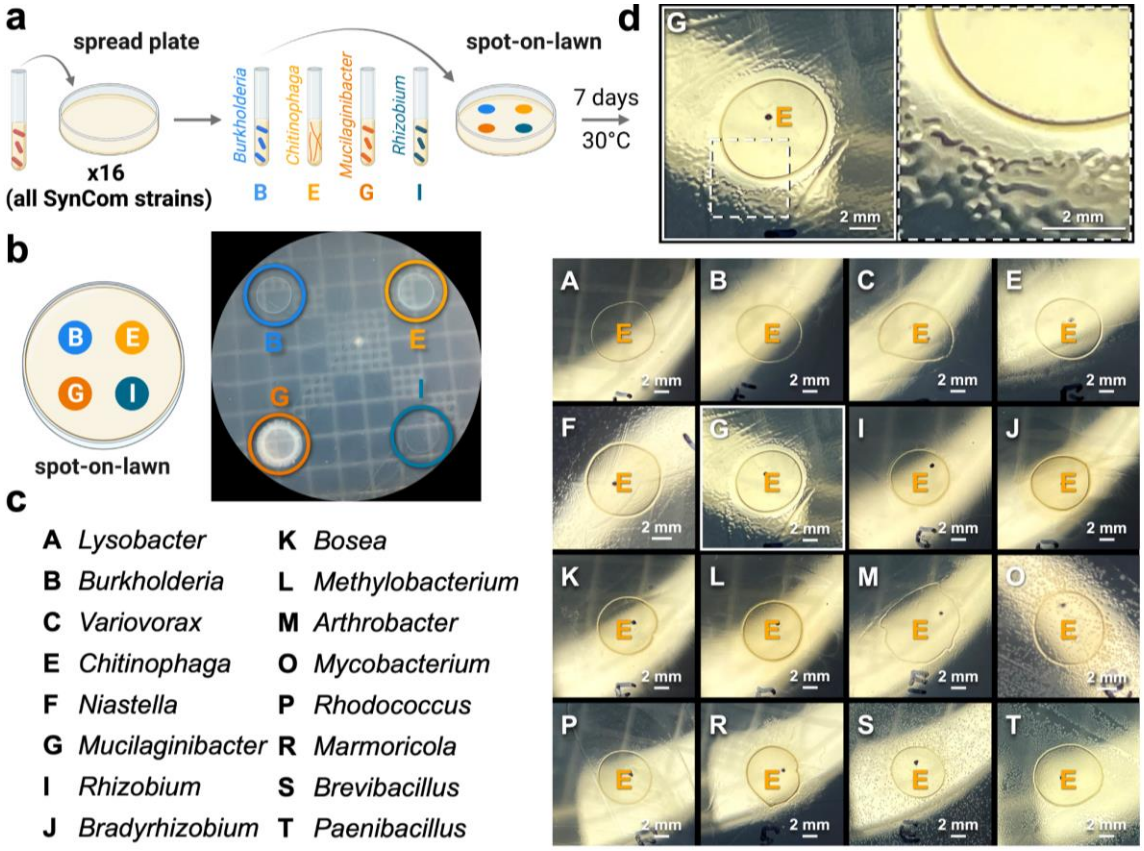


**Suppl. Fig. S8. Guild-based predictions of competitive interactions reveal guild-specific antimicrobials. a)** Syncom members from strong competition pairs identified by guild classification were spotted onto lawns of the 16 SynCom members; **b)** representative image of spot-on-lawn plates after 6-9 days of incubation; **c)** 16 SynCom members key; **d)** (bottom) representative images of *Chitinophaga* spot on lawns of the 16 SynCom members, with the *Chitinophaga*-*Mucilaginibacter* competition pair highlighted. (top) *Chitinophaga*-*Mucilaginibacter* competition pair with white box to emphasize the zone of inhibition around *Chitinophaga* spot.

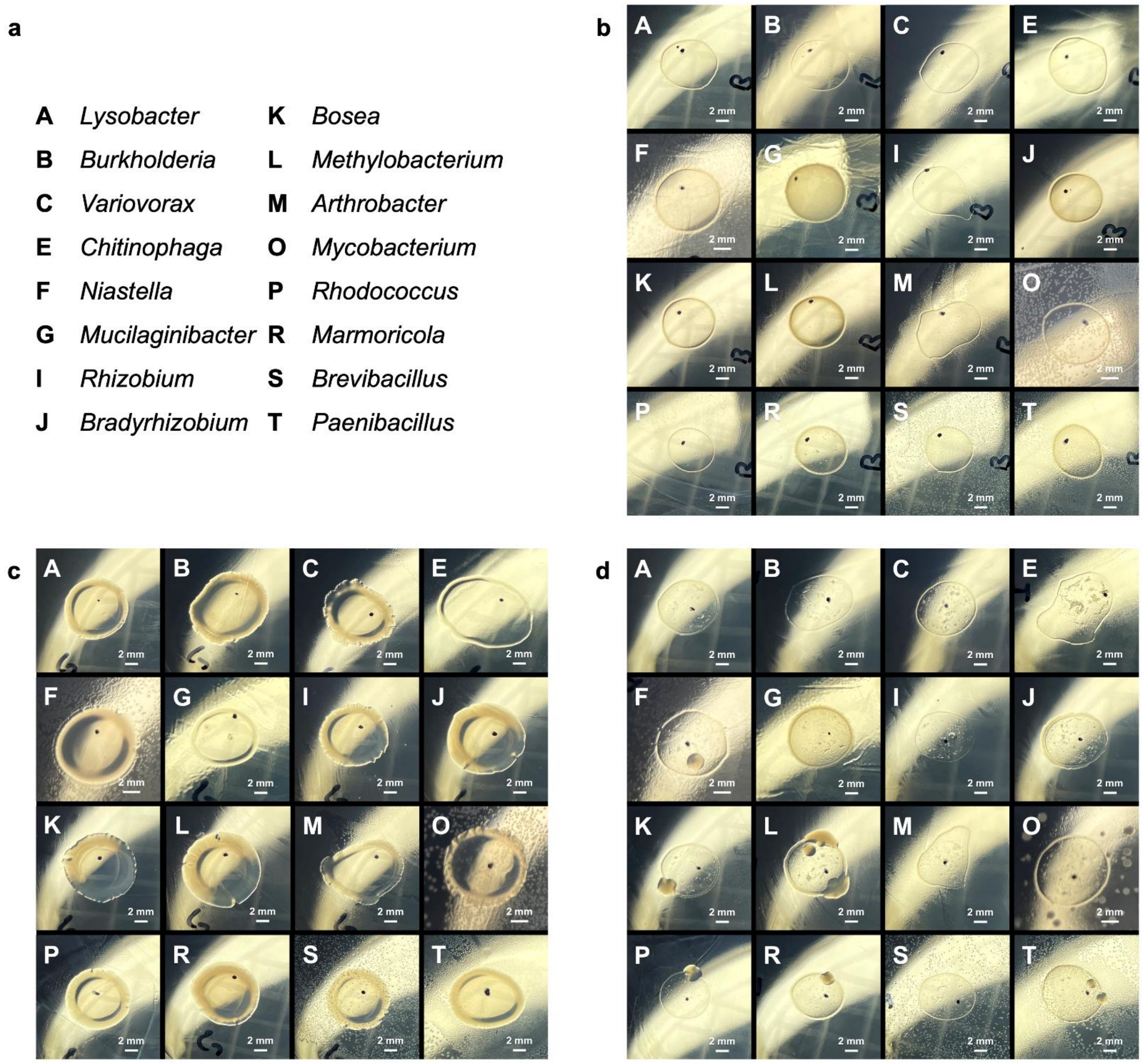


**Suppl. Fig. S9. Spot-on-lawn assays of competition pairs predicted by guild classification.** **a)** 16 SynCom member key; **b-d)** representative images of **b)** *Burkholderia*, **c)** *Mucilaginibacter*, and **d)** *Rhizobium* spot on lawns of the 16 SynCom members.

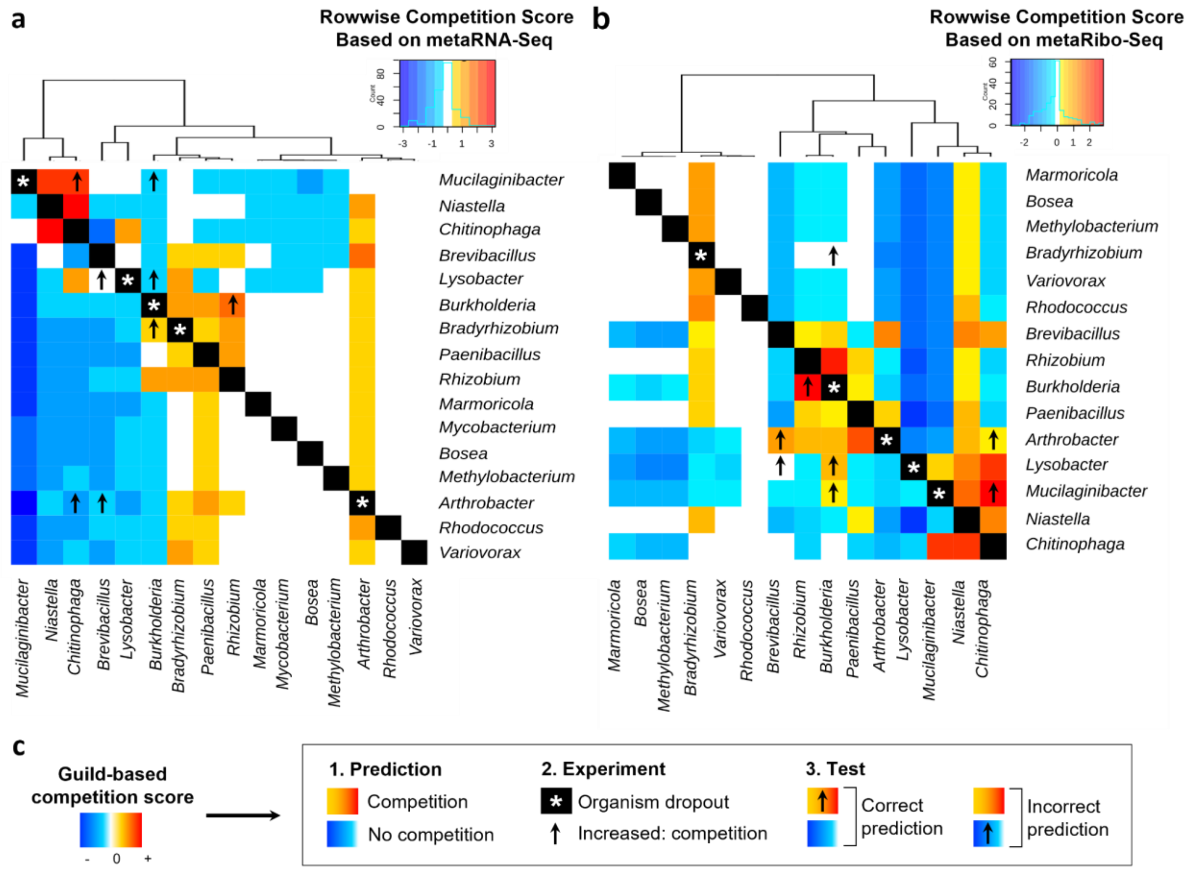


**Suppl. Fig. S10. Metatranscriptomics does not accurately predict competitive interactions in the SynCom. a)** metaRNA-Seq (metatranscriptomic) and **b)** metaRibo-Seq (metratranslatomic) guild-based competition score heatmaps showing competition score between each bacterium and the others, rowwise (see methods); **c)** Summary of the workflow used to measure prediction performance of predicted competition. A competition score was computed based on proximity of the metabolic guilds (see methods). Competition score validation was performed as follows: 1. Prediction: positive competition scores predict that the corresponding bacterium is likely to be a competitor of the reference bacterium (in row); 2. Experimental validation was performed by experimentally removing single SynCom members (asterisks) and identifying which other members increased in abundance in response to the removal (arrows); 3. Comparison of prediction with experimental results: we predicted that competitors would increase in abundance upon dropout (true positive), while non-competitors would not (true negatives). Sensitivity and specificity of prediction calculations included all comparisons for the 5 tested dropout experiments (5 rows containing an asterisk). Overall, metaRNA-Seq data poorly predicted competitive interactions (sensitivity = 37.5%, specificity = 76%, **a)**) as compared to metaRibo-Seq (sensitivity = 75%, specificity = 81%, **b)**) and TE data (sensitivity = 100%, specificity = 74%, Fig. 2d-e) emphasizing the importance of integrating translational regulation to predict microbial interactions.

**Suppl. Fig. S11. Multi-omics analysis of genes coding for metabolite import proteins.** Heatmaps showing metagenomic (left), metatranscriptomic (middle) and metatranslatomic (right) signal (row Z-score) for all genes whose names contained either “import” or “uptake”. The data show how post-genomic analyses (i.e. RNA-Seq, Ribo-Seq) considerably decrease the noise produced by genes that are not transcribed and/or translated, allowing for efficient reduction in dimensionality and for efficient design of metabolite additions. TE-based guild clustering (Fig. 1c) is shown on top of the heatmaps.

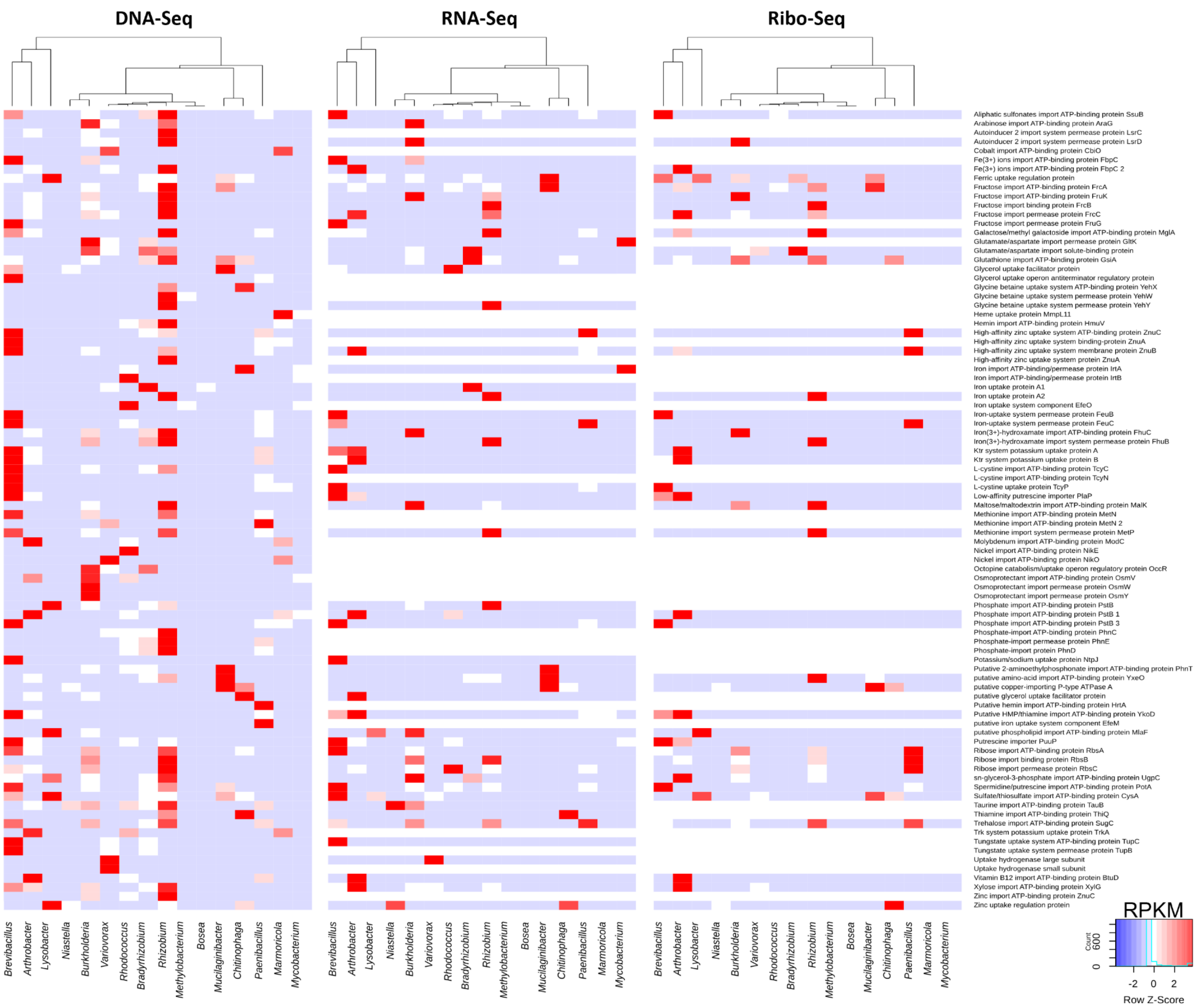

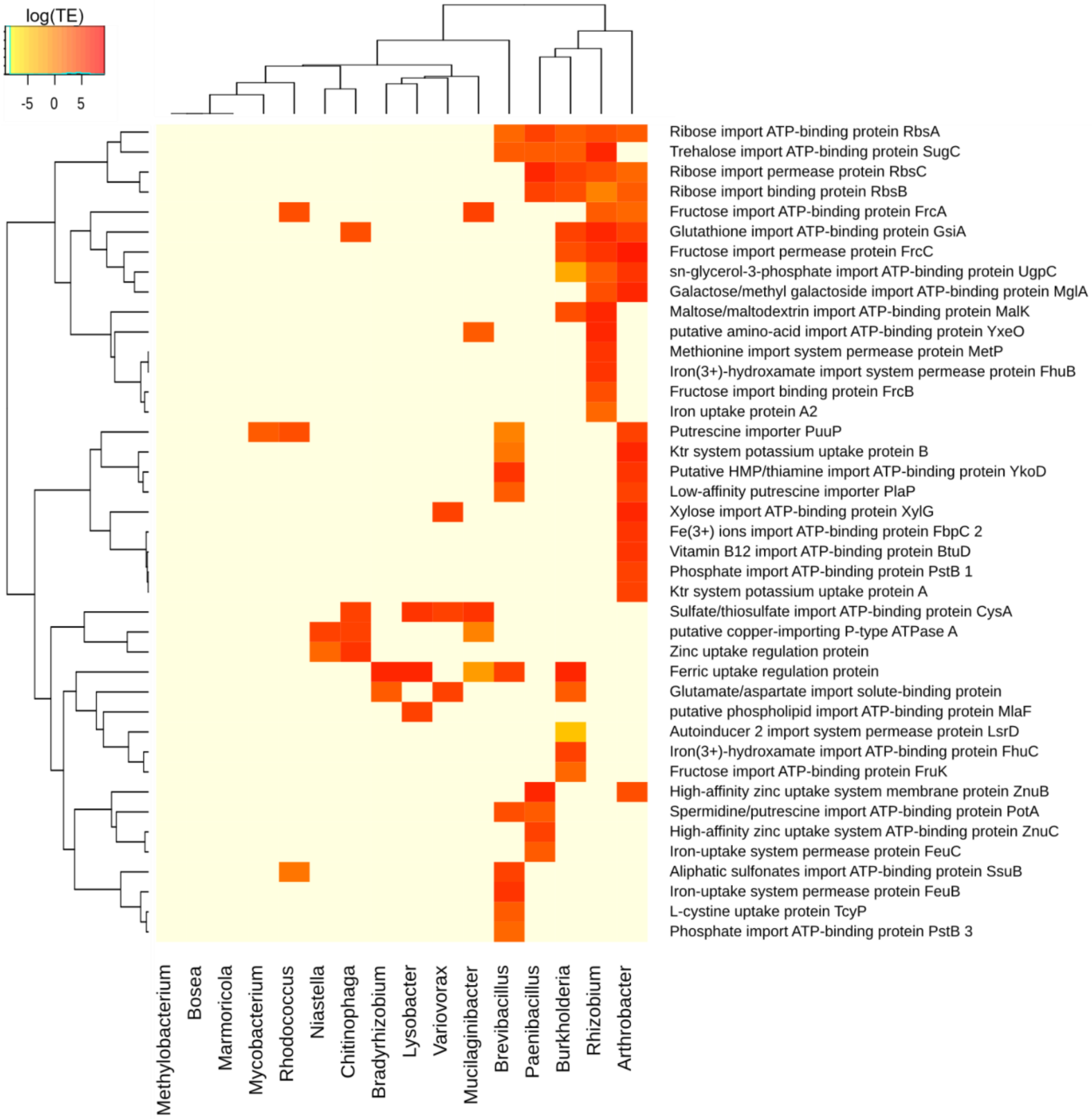


**Suppl. Fig. S12. TE for genes coding metabolite import proteins in the SynCom (log-transformed).**

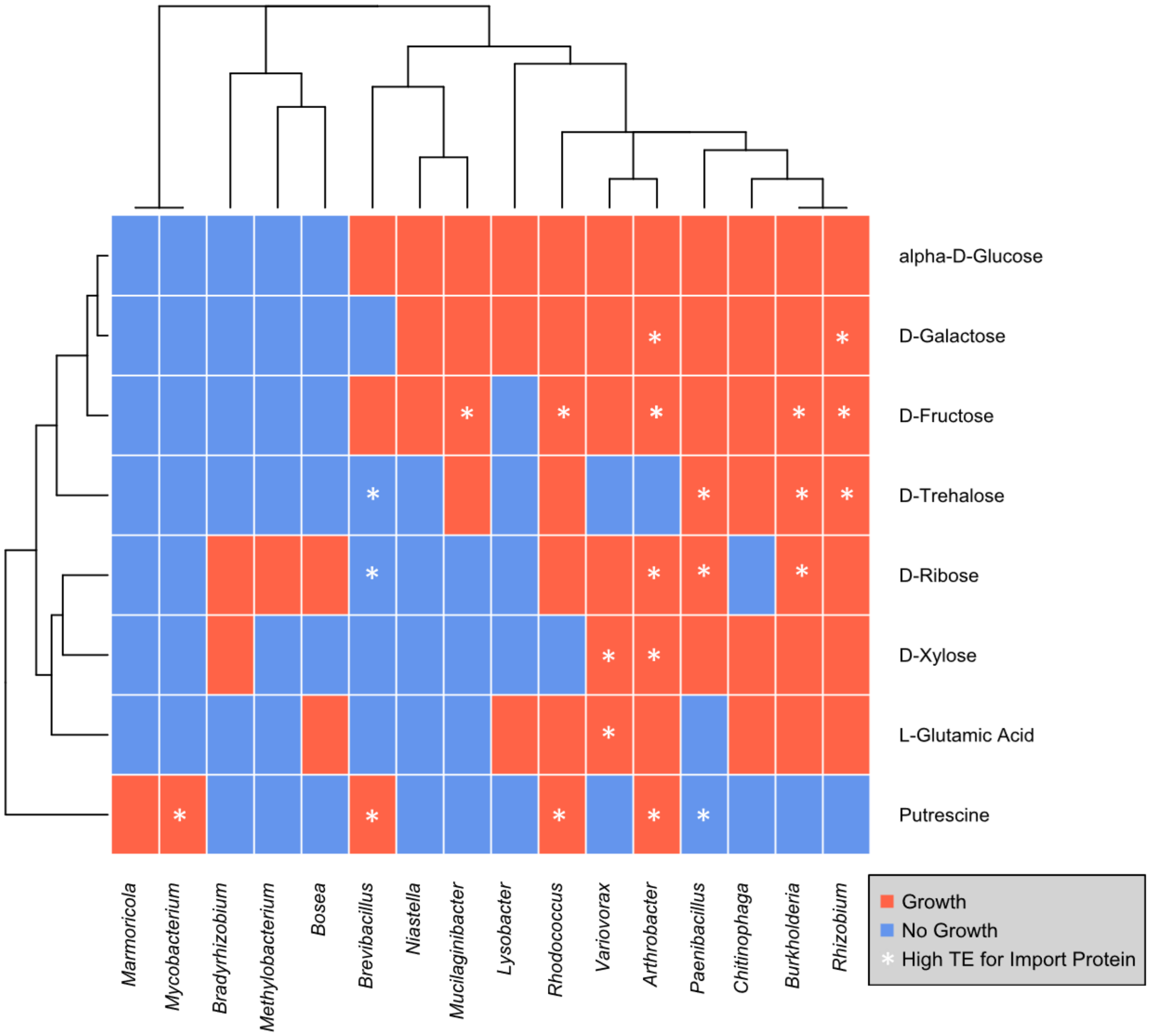


**Suppl. Fig S13. Axenic growth on substrates used as prebiotics in the SynCom.** The ability of the 16 SynCom members to metabolize specific substrates was tested using Biolog's phenotypic microarrays (see methods). Red/blue color indicate microorganisms that were/were not able to grow with the tested substrate as sole source of carbon in minimal medium. Substrates for which members exhibited a high TE for corresponding import protein(s) are indicated by white asterisks (Suppl. Fig. S12).

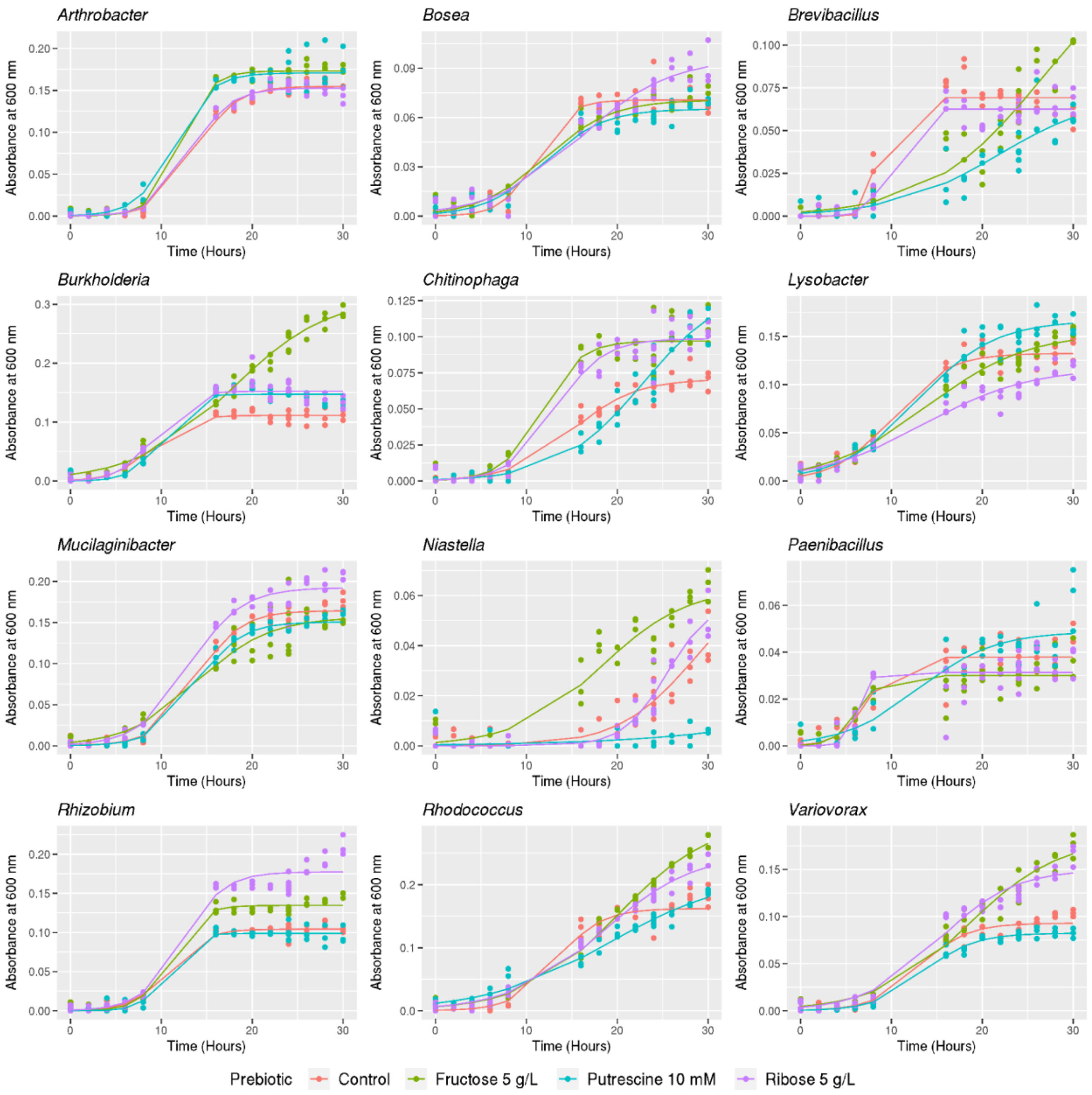


**Suppl. Fig. S14. 30-hours growth curves of SynCom isolates grown axenically at 30 °C in 0.1x R2A.** Axenic growth curves confirmed the ability of SynCom isolates to metabolize MiND-predicted preferred substrates (details in Suppl. Table S3). Note: *Bradyrhizobium*, *Marmoricola*, *Methylobacterium*, and *Mycobacterium* did not grow axenically within 30 hours in 0.1x R2A and thus are absent from the figure.

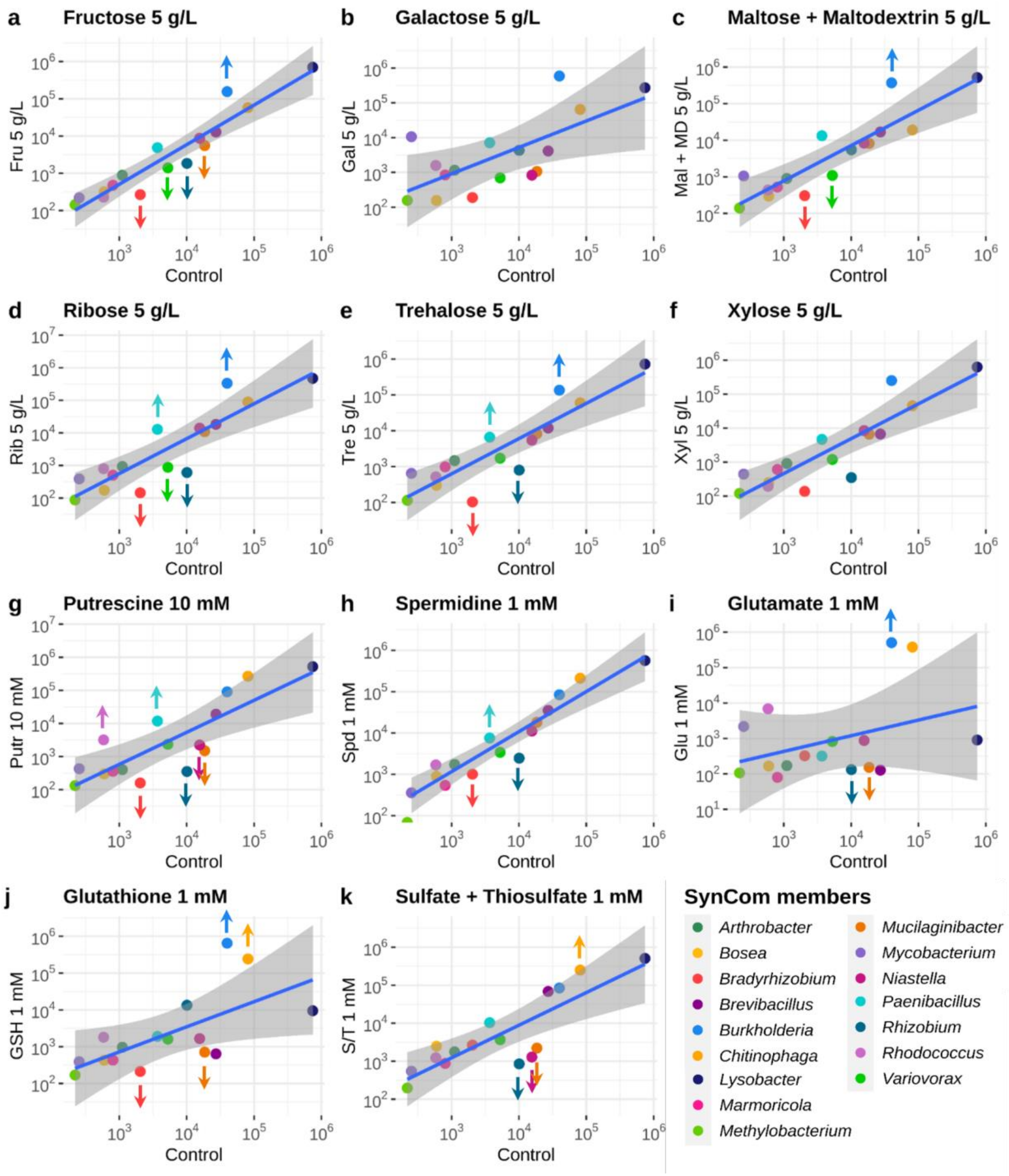


**Suppl. Fig. S15. Effects of metabolite addition on SynCom composition.** Linear regression and 99% CI of relative abundances (RPKM, log scaled) in 0.1x R2A control (x axis, n = 2) versus 0.1x R2A + substrate (y axis, n = 2). Organisms above or below the 99% CI are considered as significantly increased or decreased upon addition of substrate. Upwards arrows indicate organisms with a high TE for import protein for the tested substrate (primary targets, see Suppl. Fig. S12) that were successfully increased. Downwards arrows indicate competitors that were successfully decreased (secondary targets, see Fig. 2d). Addition of **a)** fructose, **b)** galactose, **c)** maltose + maltodextrin, **d)** ribose, **e)** trehalose, **f)** xylose, **g)** putrescine, **h)** spermidine, **i)** glutamate, **j)** glutathione, **k)** sulfate + thiosulfate. Data from the most effective of the tested concentrations are shown, all other tested concentrations are available in Suppl. Fig S16. Overall, we observed 53 significant changes in relative abundance, out of which 25% were predicted by substrate preferences (i.e. MiND), 57% by guild association (i.e. competition), and 19% remained unexplained.

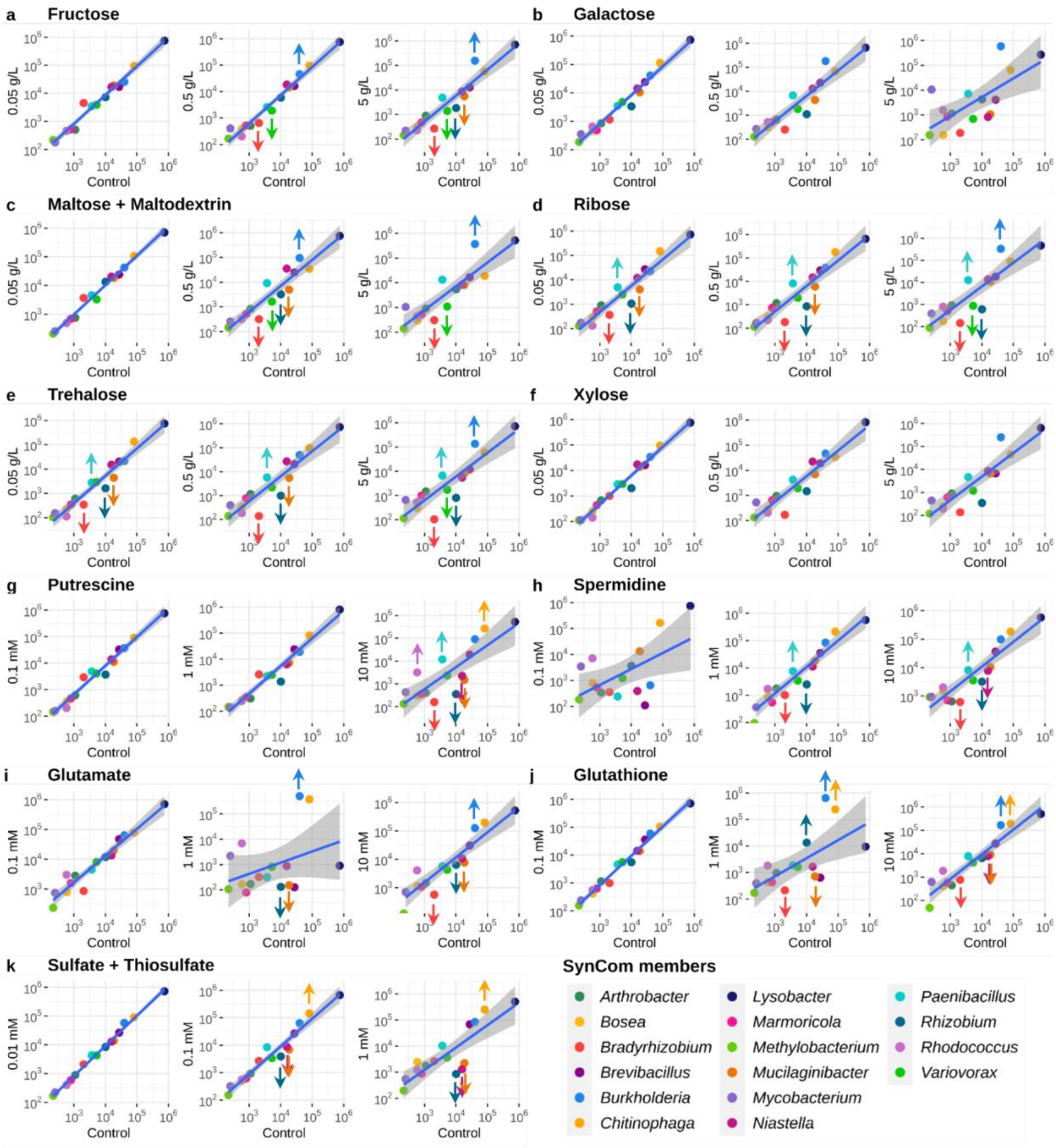


**Suppl. Fig. S16. Effects of increasing concentrations of substrate addition on SynCom composition.** Linear regression and 99% CI of relative abundances (RPKM, log scaled) in 0.1x R2A control (x axis, n = 2) versus 0.1x R2A + substrate (y axis, n = 2) for 3 tested concentrations of substrates. Organisms above or below the 99% CI are considered as significantly increased or decreased upon addition of substrate. Upwards arrows indicate organisms with a high TE for import protein for the tested substrate (primary targets, see Suppl. Fig. S12) that were successfully increased. Downwards arrows indicate competitors that were successfully decreased (secondary targets, see Fig. 2d). Addition of **a)** fructose, **b)** galactose, **c)** maltose + maltodextrin, **d)** ribose, **e)** trehalose, **f)** xylose, **g)** putrescine, **h)** spermidine, **i)** glutamate, **j)** glutathione, **k)** sulfate + thiosulfate.

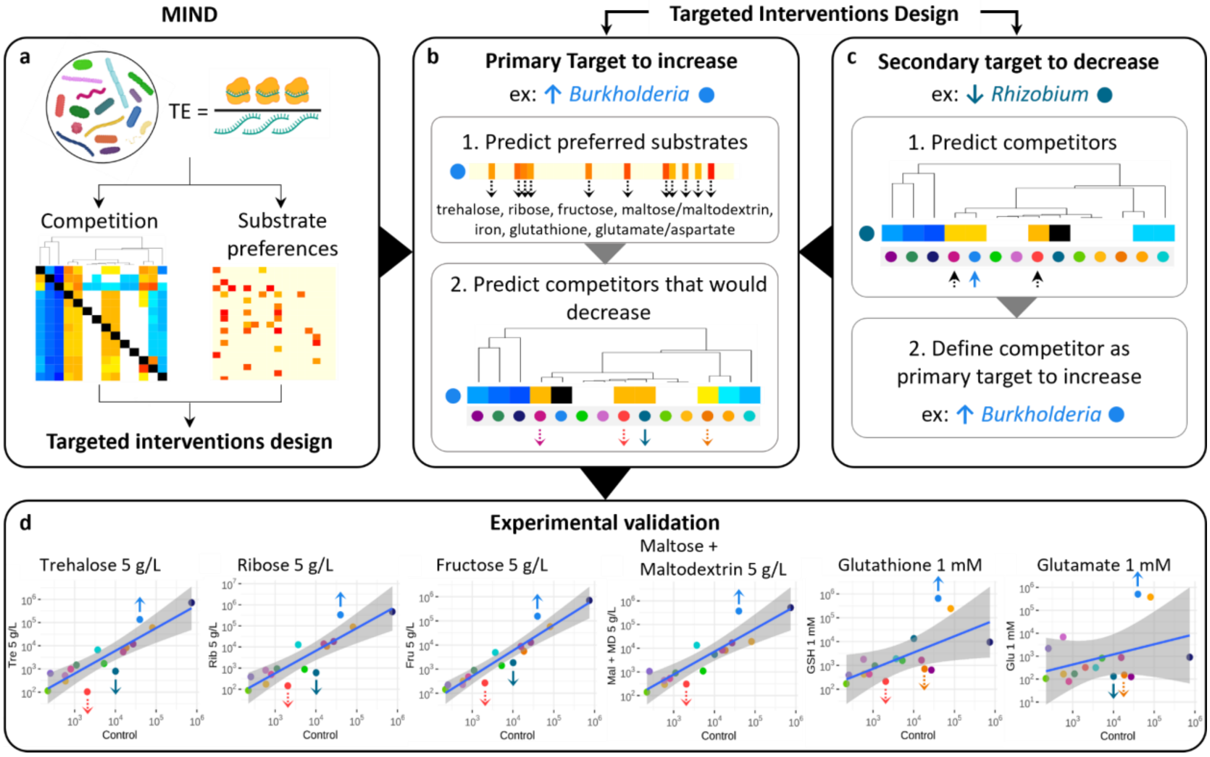


**Suppl. Fig. S16. Flowchart and illustrative examples of MiND to design targeted interventions in a microbial community. a)** MiND is based on TE, computed as the ratio between metatranslatomic and metatranscriptomic signals. It predicts competition and substrates preferences, and allows the design of targeted interventions in a microbial community (see details Fig. 2, 3 and Suppl. Fig S12); **b-c)** Here we distinguish two types of targeted interventions, whether the main objective is to increase (**b)**) or decrease (**c)**) the relative abundance of one specific bacterium; **b)** To increase the relative abundance of one primary target (e.g. *Burkholderia*), the workflow is 1. predict primary target's preferred substrates based on TE on import proteins (Suppl. Fig. S8); 2. predict competitors that might decrease if primary target is successfully increased in abundance (Fig 2c); **c)** To decrease the relative abundance of one specific secondary target (e.g. *Rhizobium*), the workflow is 1. predict secondary target's competitors (Fig 2c); 2. define one competitor as primary target to increase (e.g. *Burkholderia*); then follow workflow **b)** to increase this primary target; **d)** experimental validation of prebiotic interventions designed to increase primary target *Burkholderia* and decrease secondary target *Rhizobium* in the SynCom (extracted from Suppl. Fig. S15). Note, prebiotic intervention depicted in b) 1. can be replaced by or coupled with a probiotic intervention (i.e. primary target addition).

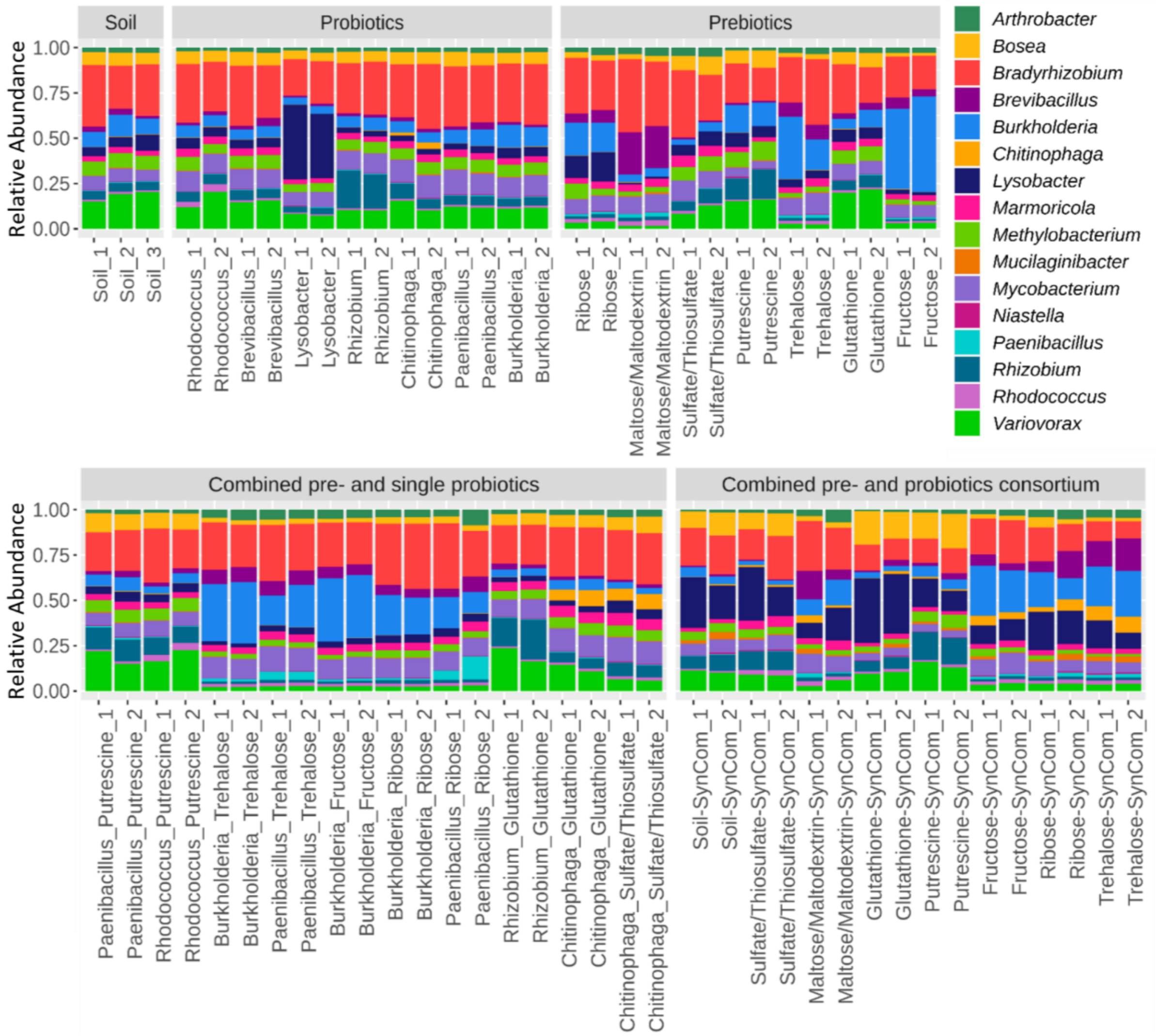


**Suppl. Fig. S18. Overview of relative abundances after pre- and probiotics interventions in soil.** Relative abundances of SynCom members grown together in soil after 7 days of growth at 30 °C for all the tested conditions (2-3 replicates each).

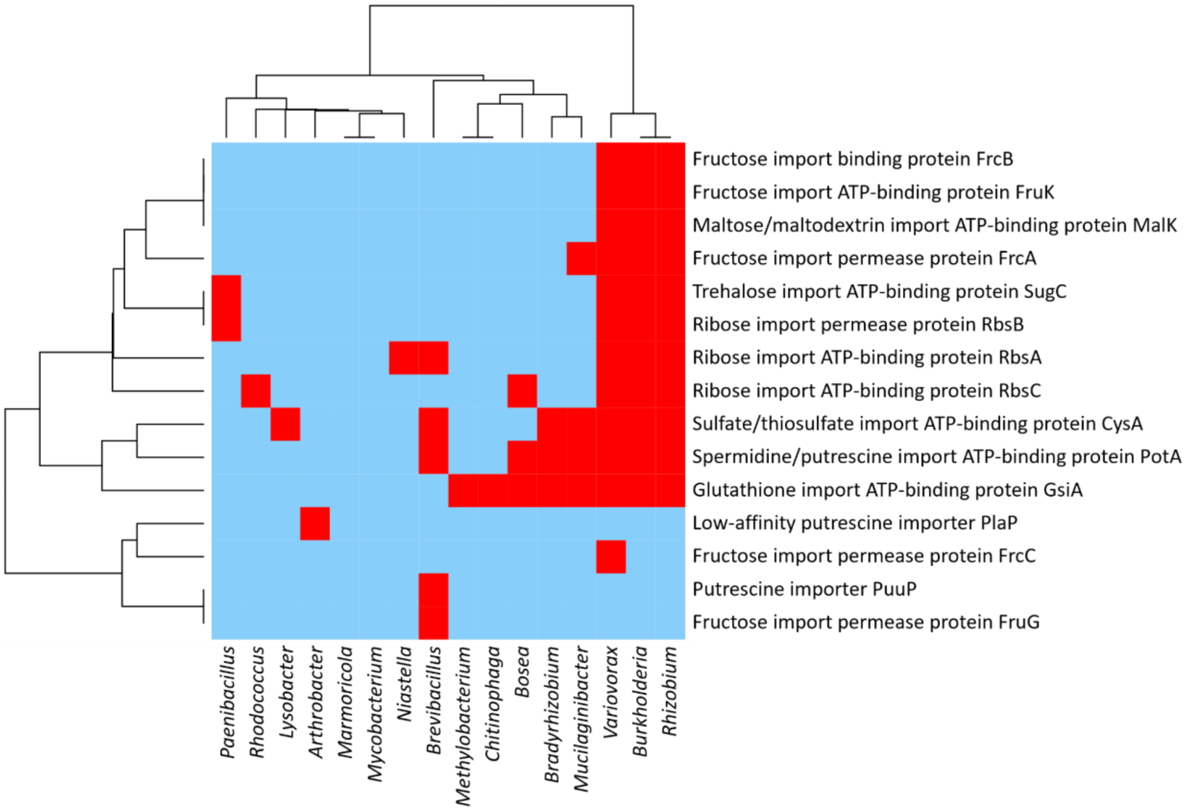


**Suppl. Fig. S19. MiND in a 16 member SynCom grown together with soil (tested prebiotics only).** Detection of transcriptional and/or translational activity (red) for a given substrate's import protein defines microbial niches. Only import proteins for the tested substrates are shown.

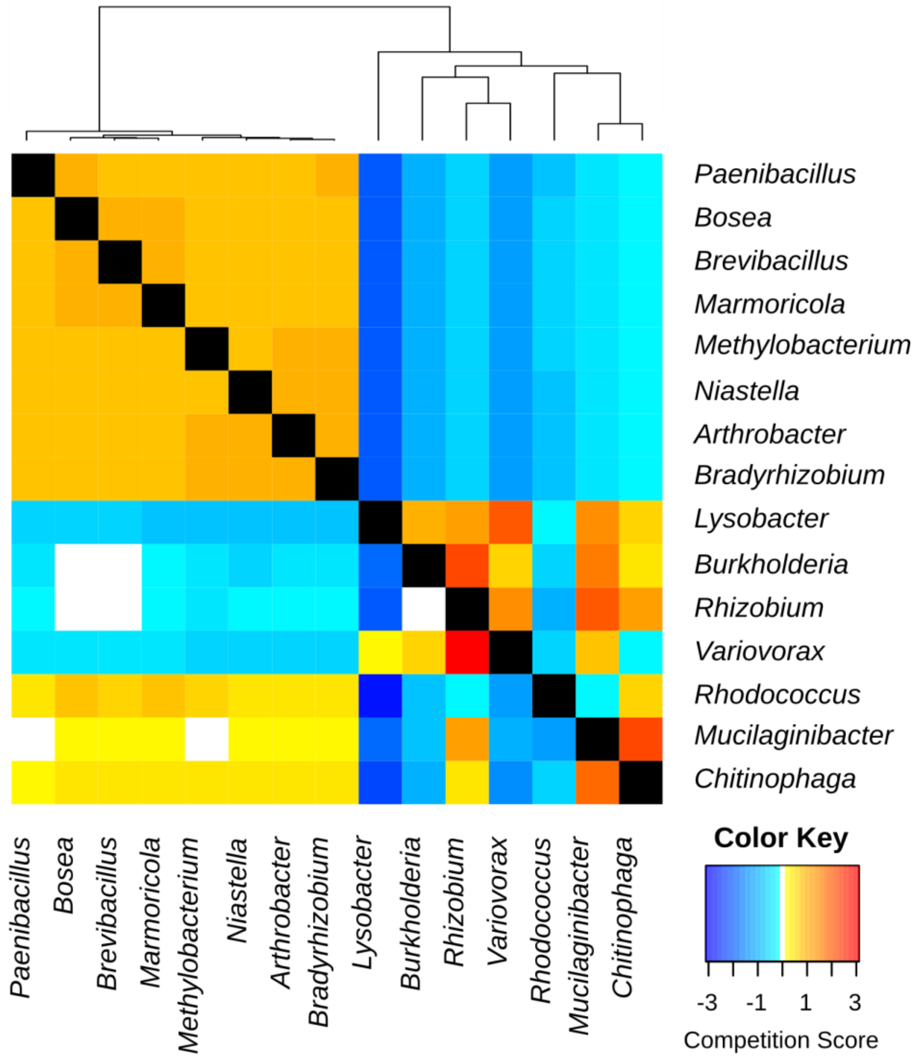


**Suppl. Fig. S20. Guild clustering (top) and competition score based on TE measured in 16 SynCom members grown together in soil.**

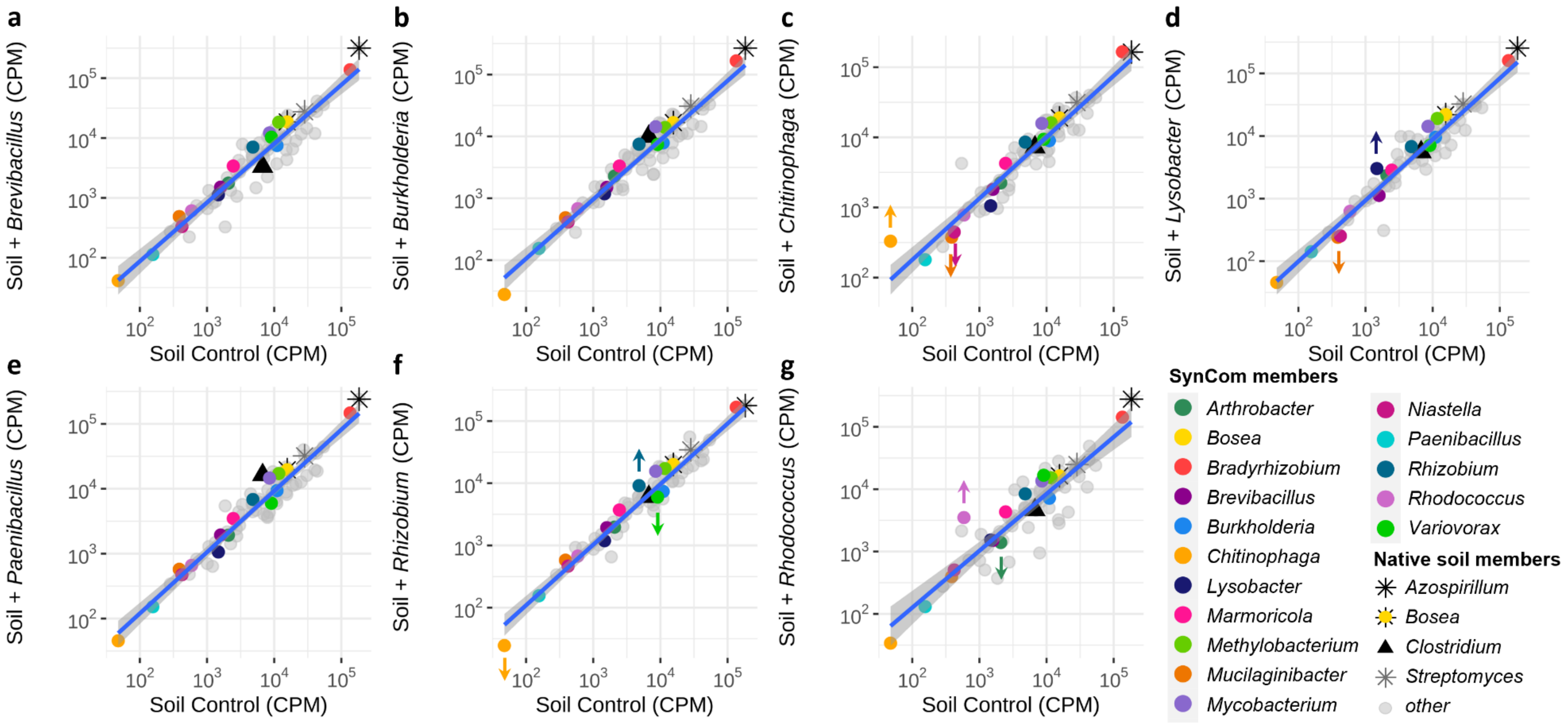


**Suppl. Fig. S21. Effects of probiotic intervention in soil.** Linear regression and 99% CI of metagenomics relative abundances (CPM, log scaled) in soil grown in 0.1x R2A control (x axis) versus 0.1x R2A + probiotic (y axis). Organisms above or below the 99% CI are considered as significantly increased or decreased upon addition of substrate. Upwards arrows indicate probiotic primary targets. Addition of **a)** *Burkholderia*, **b)** *Brevibacillus*, **c)** *Chitinophaga*, **d)** *Lysobacter*, **e)** *Paenibacillus*, **f)** *Rhizobium*, and **g)** *Rhodococcus*.

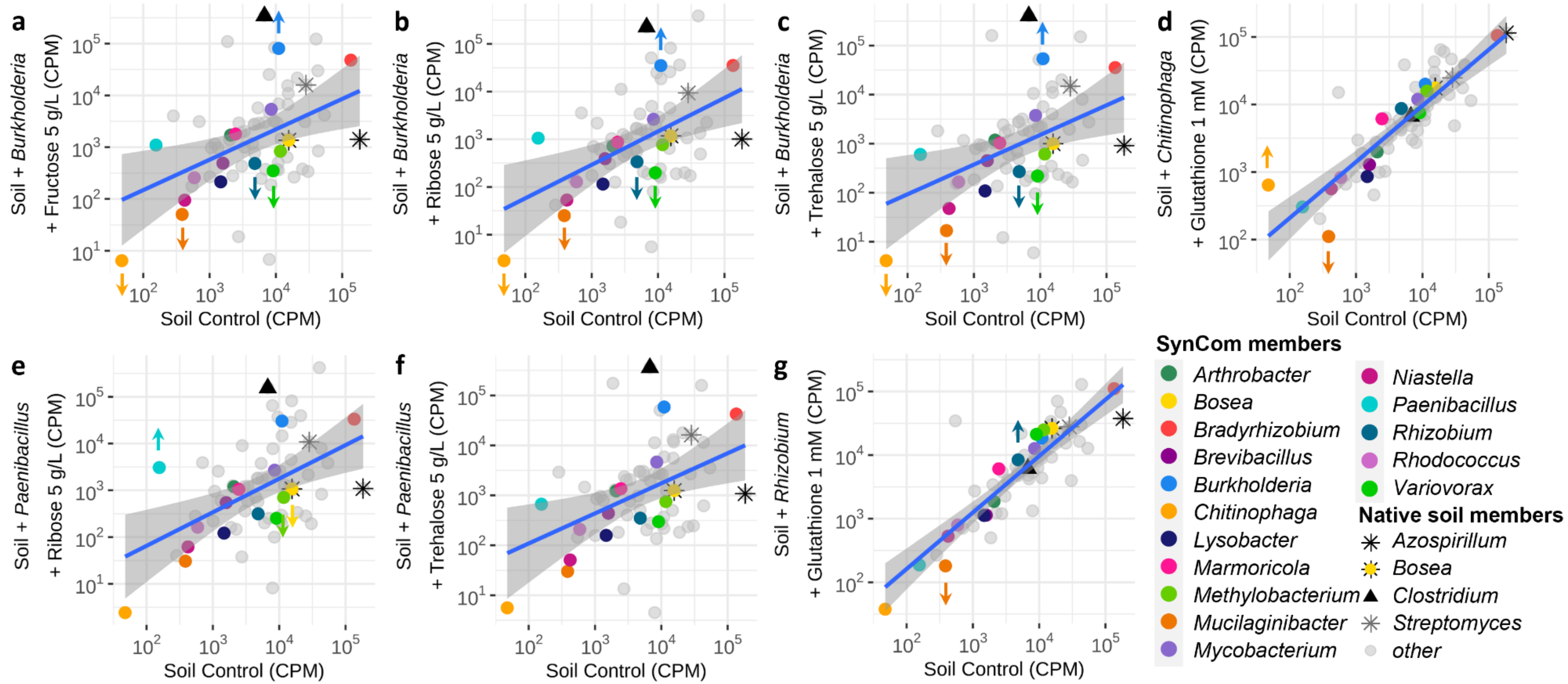


**Suppl. Fig. S22. Effects of combined niche prebiotic and single probiotic intervention in soil.** Linear regression and 99% CI of metagenomics relative abundances (CPM, log scaled) in soil (x axis) versus soil + single probiotic + prebiotic (y axis). Organisms above or below the 99% CI are considered as significantly increased or decreased upon addition of substrate. Upwards arrows indicate successfully increased primary targets, downward arrows indicate successfully decreased secondary targets. Addition of **a)** *Burkholderia* + fructose 5 g/L, **b)** *Burkholderia* + ribose 5 g/L, **c)** *Burkholderia* + trehalose 5 g/L, **d)** *Chitinophaga* + glutathione 1 mM, **e)** *Paenibacillus* + ribose 5 g/L, **f)** *Paenibacillus* + trehalose 5 g/L, and **g)** *Rhizobium* + glutathione 1 mM.

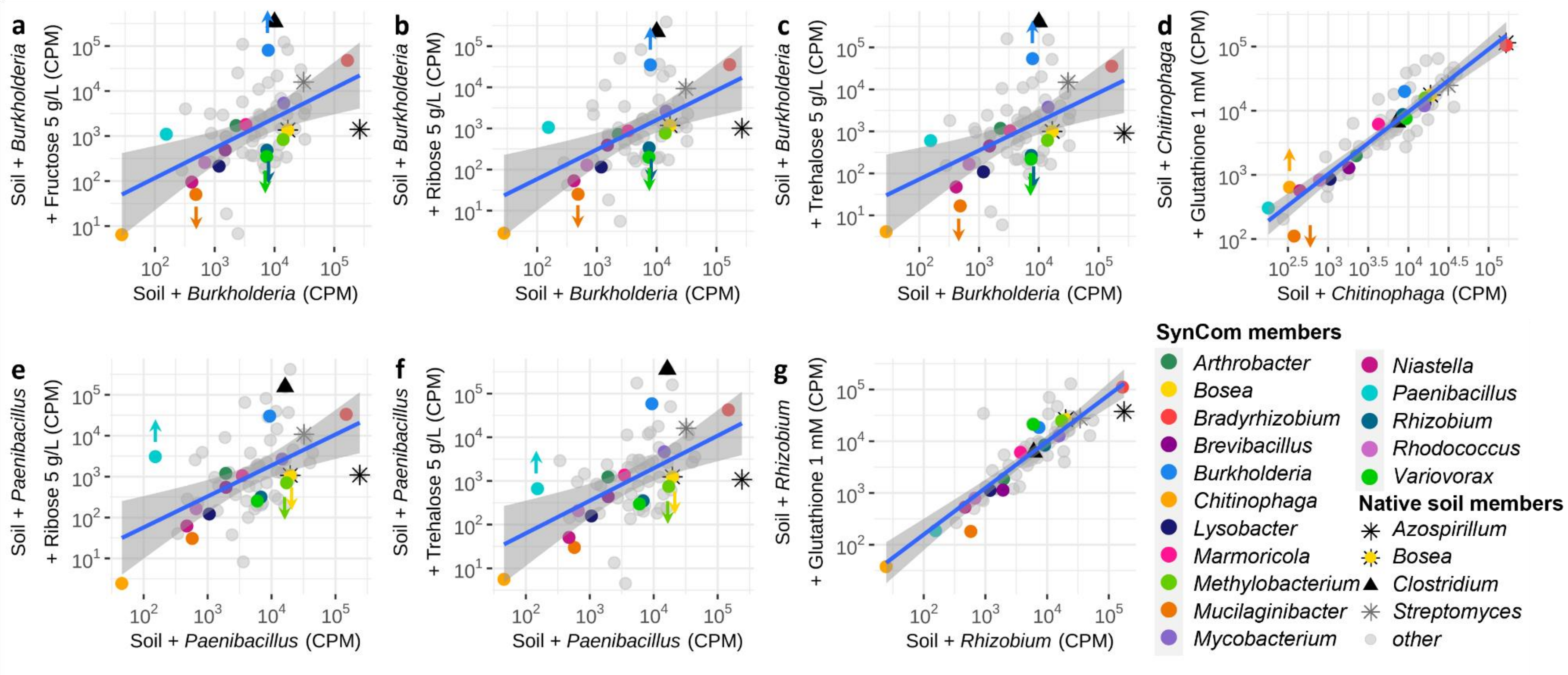


**Suppl. Fig. S23. Effect of combined niche prebiotic and single probiotic intervention exceeds the effect of probiotic treatment alone in soil.** Linear regression and 99% CI of metagenomics relative abundances (CPM, log scaled) in soil + single probiotic (x axis) versus soil + single probiotic + prebiotic (y axis). Organisms above or below the 99% CI are considered as significantly increased or decreased upon addition of substrate. Upwards arrows indicate successfully increased primary targets, downward arrows indicate successfully decreased secondary targets. Addition of **a)** *Burkholderia* + fructose 5 g/L, **b)** *Burkholderia* + ribose 5 g/L, **c)** *Burkholderia* + trehalose 5 g/L, **d)** *Chitinophaga* + glutathione 1 mM, **e)** *Paenibacillus* + ribose 5 g/L, **f)** *Paenibacillus* + trehalose 5 g/L, and **g)** *Rhizobium* + glutathione 1 mM.

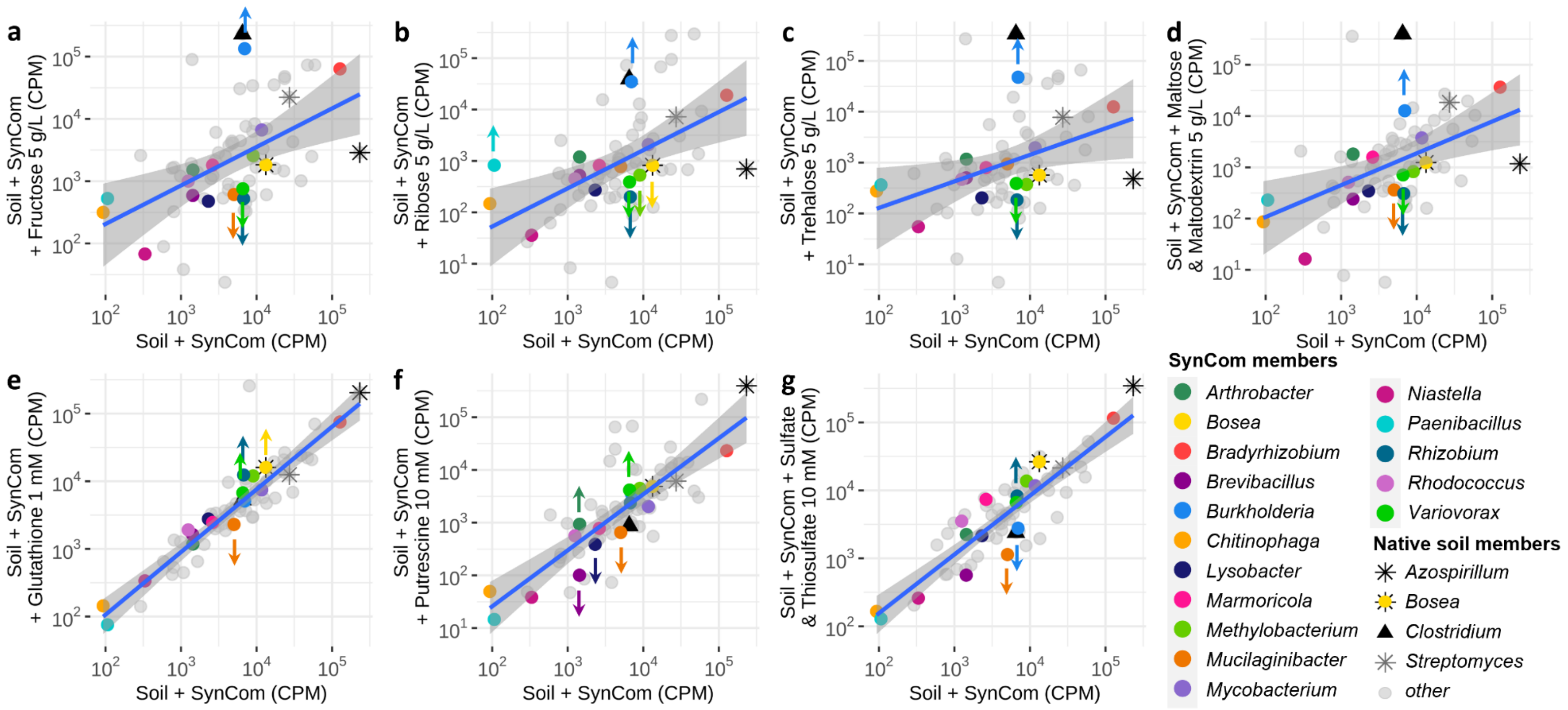


**Suppl. Fig. S24. Effects of combined pre- and probiotic consortium intervention in soil.** Linear regression and 99% CI of metagenomics relative abundances (CPM, log scaled) in soil + SynCom probiotic (x axis) versus soil + SynCom probiotic + prebiotic (y axis). Organisms above or below the 99% CI are considered as significantly increased or decreased upon addition of substrate. Upwards arrows indicate successfully increased primary targets, downward arrows indicate successfully decreased secondary targets. Addition of **a)** fructose 5 g/L, **b)** ribose 5 g/L, **c)** trehalose 5 g/L, **d)** maltose + maltodextrin 5 g/L, **e)** glutathione 1 mM, **f)** putrescine 10 mM,and **g)** sulfate + thiosulfate 10 mM.

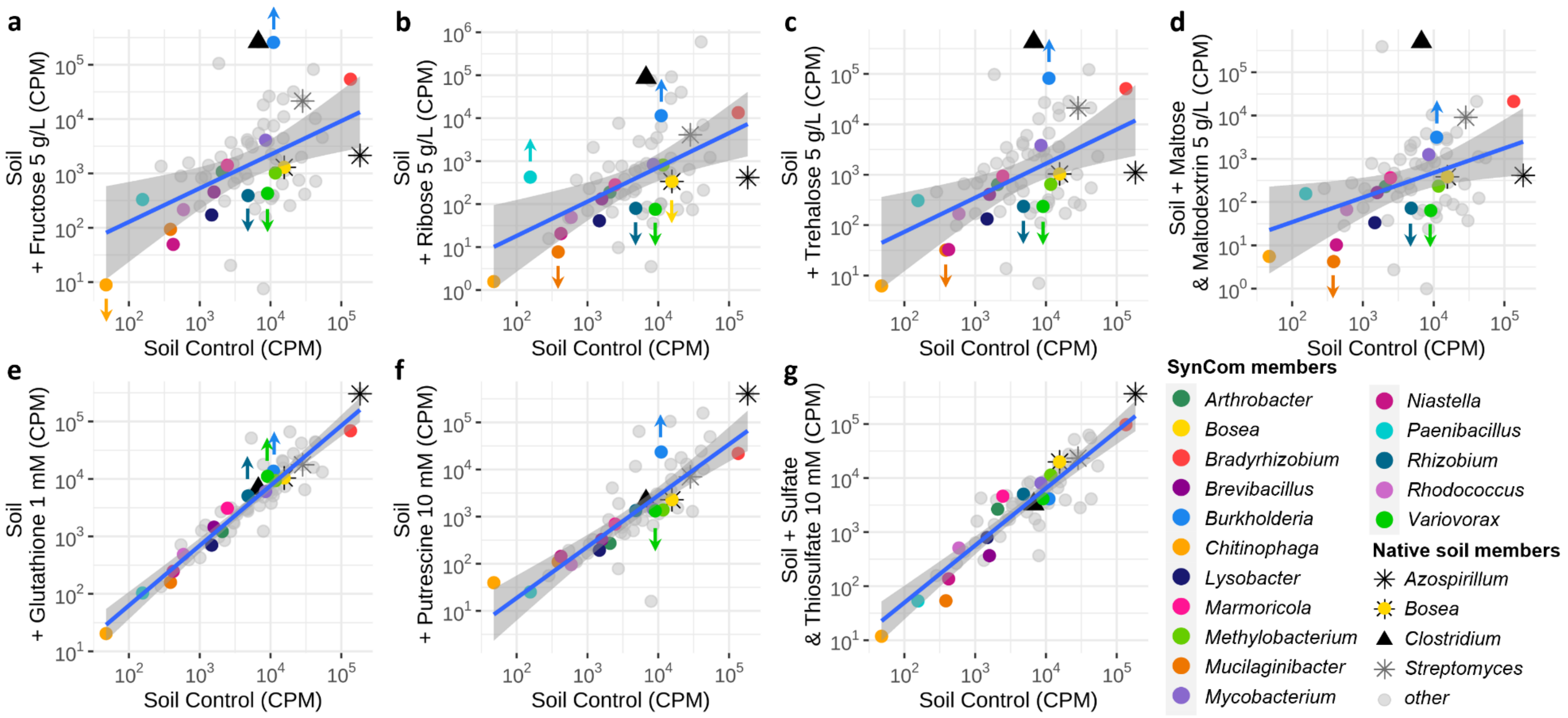


**Suppl. Fig. S25. Effects of prebiotic intervention in soil.** Linear regression and 99% CI of metagenomics relative abundances (CPM, log scaled) in soil (x axis) versus soil prebiotic (y axis). Organisms above or below the 99% CI are considered as significantly increased or decreased upon addition of substrate. Upwards arrows indicate successfully increased primary targets, downward arrows indicate successfully decreased secondary targets. Addition of **a)** fructose 5 g/L, **b)** ribose 5 g/L, **c)** trehalose 5 g/L, **d)** maltose + maltodextrin 5 g/L, **e)** glutathione 1 mM, **f)** putrescine 10 mM, and **g)** sulfate + thiosulfate 10 mM.

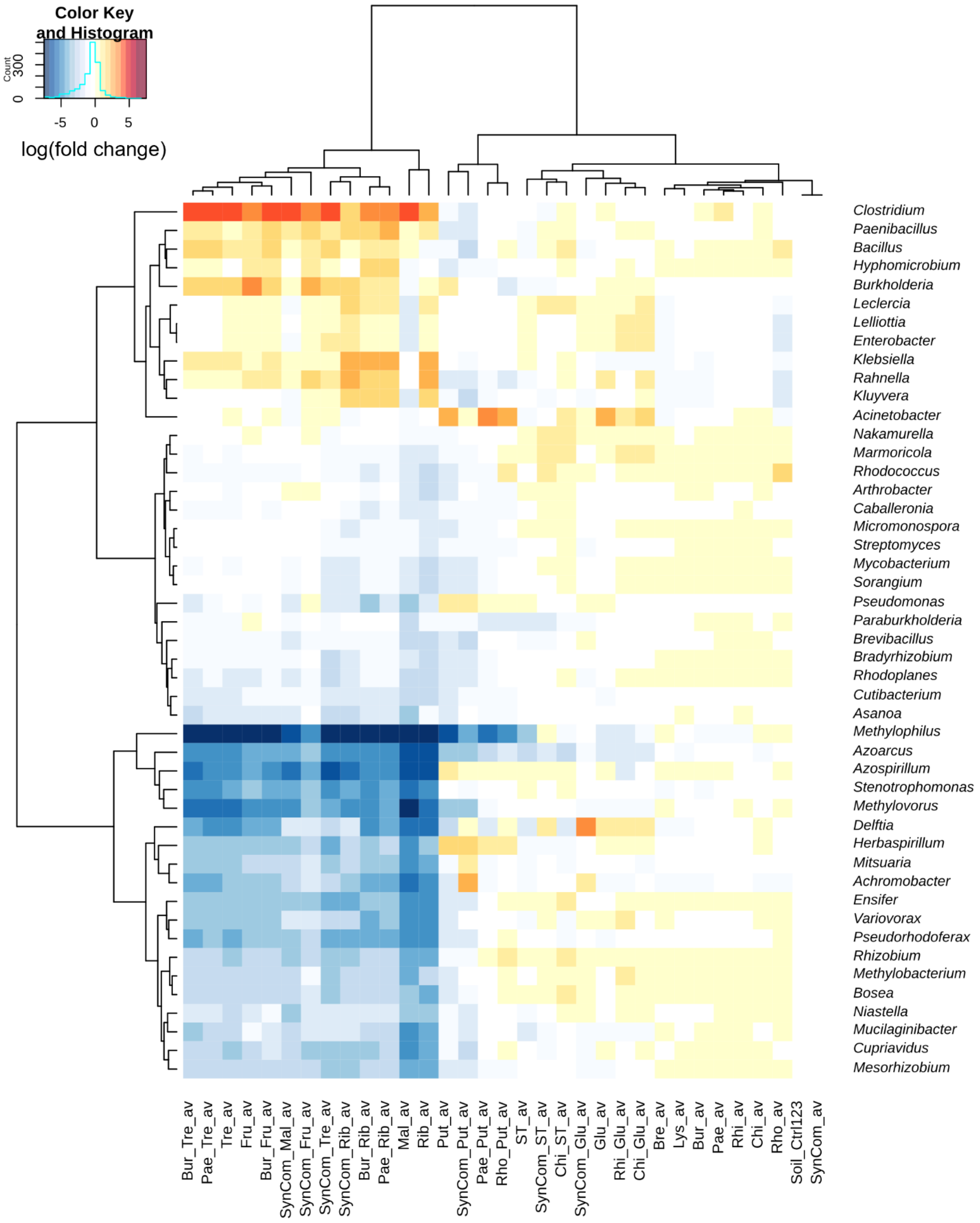


**Suppl. Fig. S26. Relative abundance changes observed in response to 30 different pre and/or probiotics treatments applied to soil.** Heatmap of log(fold change) of each bacterium's relative abundance (CPM) in all tested conditions as compared to its control. Pre and/or single strain probiotic treatments (n = 2 each) were compared to soil alone (Soil_Ctrl123, n = 3), while SynCom + prebiotic treatments (n = 2 each) were compared to Soil + SynCom (SynCom_av, n = 2). Key: av: average; probiotics: Bre: *Brevibacillus*, Bur, *Burkholderia*, Chi: *Chitinophaga*, Lys: *Lysobacter*, Pae, *Paenibacillus*, Rhi: *Rhizobium*, Rho: *Rhodococcus*; SynCom: SynCom; prebiotics: Fru: fructose, Glu: glutathione, Mal: maltose + maltodextrin, Put: putrescine, Rib: ribose, ST: sulfate + thiosulfate, Tre: trehalose.

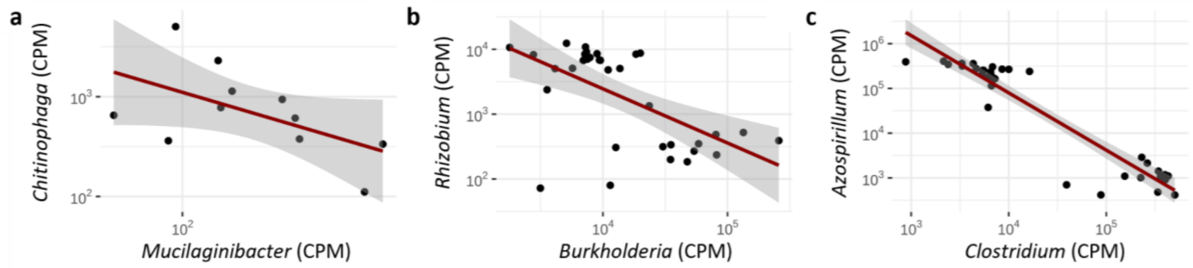


**Suppl. Fig. S27. Negative correlation between competitors observed across 30 different pre- and probiotic interventions in soil.** Relative abundances (CPM) of strong competitors **a**) *Chitinophaga-Mucilaginibacter*, **b**) *Burkholderia-Rhizobium*, and **c**) *Clostridium-Azospirillum*.

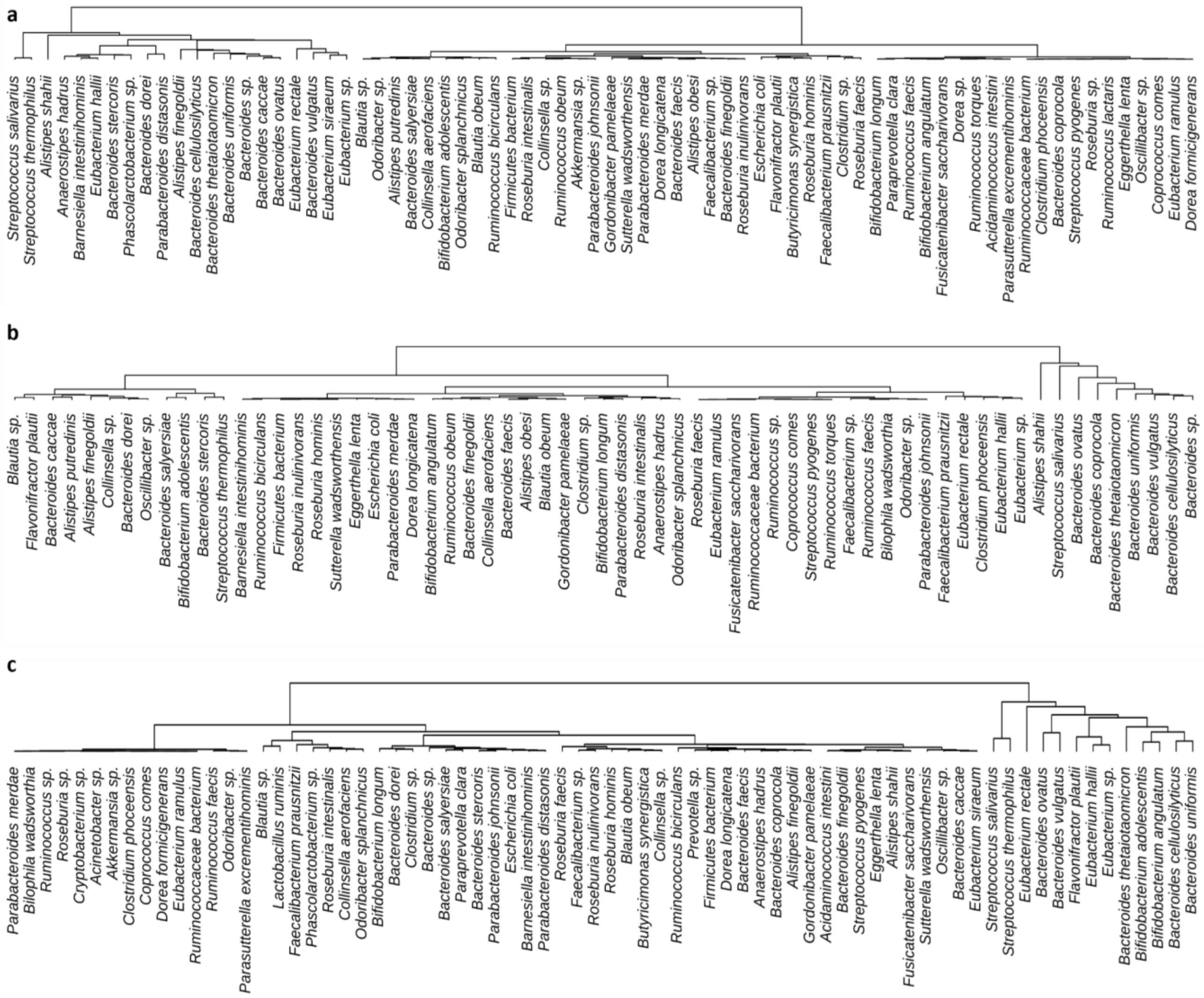


**Suppl. Fig. S28. Application of TE-based guild classification to human fecal samples. a-c)** Clustering dendrogram of TE measured on metabolic pathways in fecal samples from 3 healthy individuals (representative examples; total samples tested = 7).

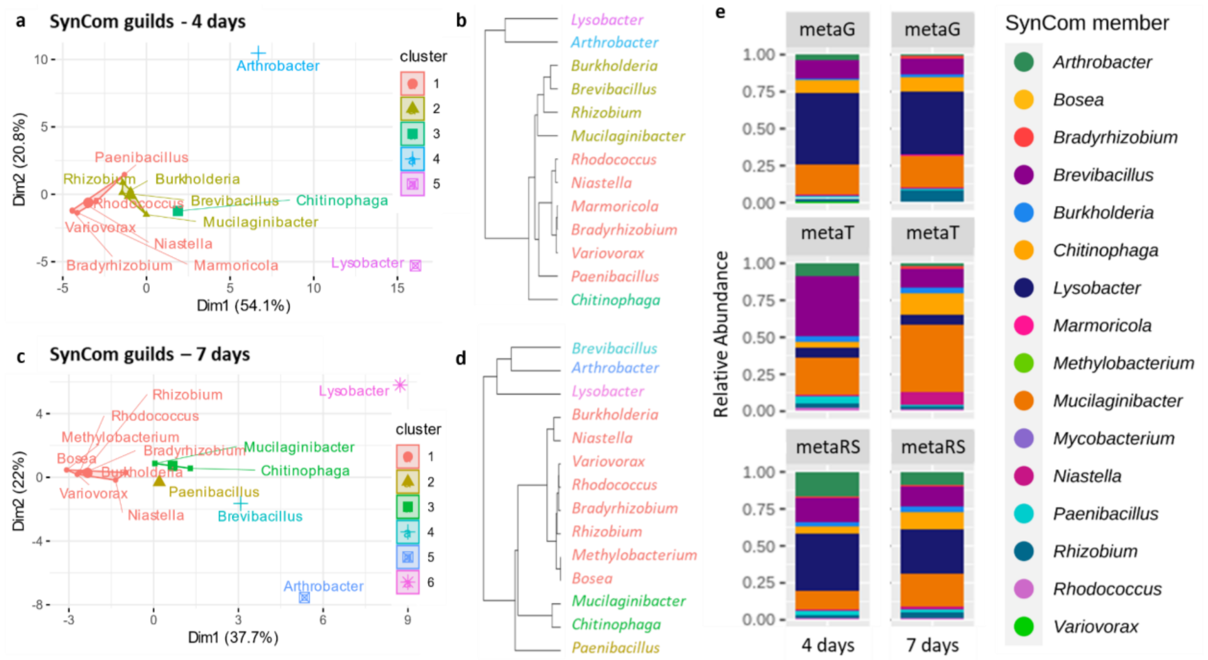


**Suppl. Fig. S29. Stability of guild-based classification of a SynCom over time. a-d)** TE-based classification of a 16-member SynCom of rhizosphere isolates after growth in complex medium for 4 (**a,b**) and 7 (**c,d**) days; **e)** multi-omic taxonomic profiles (n = 4) of the 16-member SynCom after 4 (left) and 7 (right) days of growth. From top to bottom: metagenomics (metaG), metatranscriptomics (metaT), metatranslatomics, i.e. Ribo-Seq (metaRS).

#### EXTENDED DATA TABLES

**Suppl. Table S1. Summary of KEGG pathways significantly separating TE-based guilds in a 16-member SynCom.** Output of HCPC analysis[52].

**Suppl. Table S2. Determination of isolate's ability to metabolize 275 substrates by phenotypic microarray (Biolog) for the 16 SynCom members.**

**Suppl. Table S3. MiND predictions of prebiotic interventions outcomes in the SynCom.** This table compares predictions of substrate preferences made from TE information measured directly in the SynCom (i.e. MiND), with predictions made from axenic culture assays (i.e. Biolog, growth curves). It also compiles experimental results from prebiotic supplementation experiments, and allows to calculate MiND sensitivity and specificity; TP = true positive (MiND-predicted preferred substrate, experimentally validated by primary target increase), TN = true negative (MiND-predicted non-preferred substrate, experimentally validated by no primary target increase), FP = false positive (MiND-predicted preferred substrate, but not experimentally validated by primary target increase), FN = false negative (MiND-predicted non-preferred substrate, but not experimentally validated by no primary target increase). Guild-predicted competition interactions with increased primary targets are also indicated, and experimentally validated when competitors (i.e. secondary targets) are decreased.

**Suppl. Table S4. GEMs-predicted growth rates of SynCom members on growth media with and without selected substrates.** This table compares predicted growth rates (1/h) of SynCom members while simulating their genome-scale metabolic models (GEMs) with and without inclusion of selected substrates in the medium (see methods). The exchange reactions serve as the link between the external substrates provided in the medium and the internal metabolic network in the model. The column labeled "Model refinement (based on BIOLOG data)" indicates whether or not the model was/was not ("yes/no") refined by incorporating growth data obtained from Biolog experiments.

#### SUPPLEMENTARY MATERIAL

**Suppl. Material 1. MetaRibosome profiling (MetaRibo-Seq) protocol used in this study.**
