## Supplementary material for "Guild and Niche Determination Enable Targeted Alteration of the Microbiome": MetaRibosome Profiling Protocol

### MetaRibo-seq protocol

#### BACTERIAL LYSIS AND PROCESSING

1. Set refrigerated centrifuge at 4°C.
2. Place the Beadbeater at 4°C.
3. Clean the bench and the pipettes with RNase AWAY.
4. Prepare RECIPE 1 and store on ice (>500 ul per sample).
5. Annotate one BashingBead tube (Zymo research s6012-50) per sample.
6. a) For fresh bacterial cultures samples (SynCom): centrifuge culture tube at 4°C, 7,000 rpm, 6 min. Discard supernatants. Resuspend in leftover volume and transfer to a 1.5 mL Eppendorf tube. Centrifuge at full 4°C, full speed, 2 min. Resuspend bacterial pellets in 500 ul RECIPE 1 and transfer into a BashingBead tube.

   b) For fresh cultured soil samples: add 500 ul of lysis buffer RECIPE 1 to a BashingBead tube (Zymo research s6012-50). Incubate on ice. Centrifuge culture tube containing soil + culture medium at 4°C, 7,000 rpm, 6 min. Discard supernatants. Scoop ~200 mg of soil + bacteria pellet and add to the BashingBead tube.

   c) For stool samples: on dry ice, cut the stool sample using a blade without thawing it such as it is ~ 200 mg in weight (about the size of a sweet pea). Add the stool sample to the BashingBead tube.
7. Secure the BashingBead tubes to the BeadBeater and lyse for 2x10 min at 4°C at max speed.
8. Centrifuge at 4°C full speed 5 min. If the lysate is not clear, centrifuge at 4°C full speed 3 min again.
9. Transfer cleared lysate to a new microcentrifuge tube.
10. Check the [$OD_{260}$] of 1:100 dilution in water using a Nanodrop. Use water to blank the instrument prior to taking the sample reading.

#### MNASE TREATMENT

11. The input to MNase treatment is normalized based on the absorbance measure taken on Step 10 to calculate the 25 $A_{260}$ units. The following formula should be used to determine the volume of cleared lysate to use in the next step:

$$\text{Volume of MNase (ul)} = (194 \times 12 \times 100 \times [OD_{260}]) / (1000 \times 25)$$

***Note: This is for MNase 500 U/ul. Multiply the result by 2.5 if using 200 U/ul (2,000 gel units/ul)***

***Note: Here we are talking about Kunitz units. 1 Kunitz unit = 10 gel units.***

Volume of lysate (ul) = 196 - Volume of MNase (ul)

*At this stage the leftover lysate can be stored at -80 °C in Trizol for DNA and RNA-Seq analysis.

12. Prepare the mixture described in RECIPE 3.

13. Add 10 ul RNase-free DNase (10U/ul)

14. Incubate the mixture for 2 hr at 25°C. (Set timer for 1h - 1h30).

15. After the mixture has been incubating for 1h - 1h30, begin the preparation of monosome recovery (steps 17 - 23).

16. Proceed to monosome recovery immediately or quench the reaction by adding 2.5 ul EGTA (500mM stock).

MONOSOME RECOVERY

**Preparation**

---

17. Change the centrifuge rotor (free swinging) and set up at 4°C.

18. Prepare 2 x 1.5 mL per sample of Polysome Buffer (RECIPE 4) and store on ice.

19. Prepare 2 ***Sephacryl S400 MicroSpin columns (Cytiva)*** for each sample. Shake Sephacryl S400 MicroSpin columns several times to resuspend the resin. Tap on the bench top to remove bubbles that may form. Vortex.

20. Clean the electronic serological pipettor (remove plastic nose piece and form seal with rubber insert, pulse trigger) with RNase AWAY.

21. Open the columns and gently push through the buffer with an electronic serological pipettor (remove plastic nose piece and form a seal with rubber insert, pulse trigger). Stop as soon as froth appears. **Alternatively**, use a vacuum with gentle aspiration to push out the buffer.

22. Equilibrate the resin by passing 1.5 mL Polysome Buffer (RECIPE 4) through each column in 3x500 µL aliquots emptying the collection tube after each wash. Remove the last wash by spinning for 4 min at 600 x g in a swinging-bucket table-top microcentrifuge at 4°C.

23. Discard the flow through and attach a new collection tube to each column.

---

24. Apply half of each MNase-treated sample to a single Sephacryl S400 column (not to

exceed 100 µL per column) and centrifuge for 2 min at 4 °C and 600 x g.

25. At this stage you will combine the samples processed using 2 Sephacryl S400 Columns. Add 700 ul Trizol to the flow-through and vortex immediately.

26. Wait 1 min.

*At this stage the lysate can be stored in Trizol at -80 °C.

27. Change the centrifuge rotor to fixed angle.

28. Add 140 ul of Chloroform. Vortex ~ 10 sec.

29. Centrifuge for 4 min at max speed, 4°C.

30. Set the centrifuge at room temperature.

31. Take 500ul of the aqueous phase without disturbing the white interface and transfer to a new reaction tube. Do not try to take the whole sample.

32. Add 1 volume of fresh 70% ethanol (usually 500 ul) and mix thoroughly by vortexing. Do not centrifuge.

33. Pipette 500 ul of sample into a ***RNeasy Mini spin column (Qiagen)*** placed in a 2 ml collection tube. Centrifuge at 8,000 x g (10,000 rpm) for 1 min, soft acceleration, at room temperature. **Take the flowthrough** and apply it to the column again. **Finally, pipet the flow-through** (which contains miRNA) into a 2 ml reaction tube. Repeat this step for the remaining volume of samples.

34. Apply the steps of the ***RNA Clean & Concentrator-5 kit (Zymo)*** Trizol clean-up protocol with the following modifications:

    34.1. Add 0.65 volume (650 ul for 1 mL) of 100% ethanol and mix by vortexing.

    34.2. Apply 900 ul of sample to the RNA Clean & Concentrator-5 kit column. Centrifuge at 8,000 x g for 1 min, soft acceleration, take the flowthrough and apply it to the column again. Repeat this step for the remaining volume of samples. Discard the flow-through.

    34.3. **DO NOT** bind with the Prep Buffer.

    34.4. Wash twice with the RNA Clean & Concentrator wash buffer. 700 ul, 30 s, max speed first wash and 500 ul, 2 min second wash.

    34.5. Discard the flow-through.

    34.6. Transfer the column in a RNase-free tube.

    34.7. Elute with **20 ul** RNase/DNase-free water. Centrifuge, take the flowthrough and apply it to the column again.

35. Measure the RNA concentration using the QuBit or Nanodrop.

*At this stage the isolated RNA can be stored at -80°C.

rRNA REMOVAL

36. Follow steps 1-4 ***Qiagen FastSelect-5S/16S/23S*** protocol. Briefly:

36.1. Use 12.5 ul of RNA sample maximum to get 100 ng - 1ug total RNA.

36.2. In a PCR tube, mix:

| **Component** | **Volume/reaction** |
|---|---|
| Total RNA sample | Variable (max. 12.5 ul) |
| FH Buffer, 10x | 1.5 ul |
| QIAseq FastSelect - 5S/16S/23S | 1 ul |
| Nuclease-free water | 15 ul q.s. |
| Total volume | 15 ul |

36.3. Incubate as follows (FastSelect protocol - skip RNA fragmentation (step 1)):

2 min at 75°C
2 min at 70°C
2 min at 65°C
2 min at 60°C
2 min at 55°C
2 min at 37°C
2 min at 25°C
Hold at 4°C

37. Follow the total RNA isolation protocol of ***RNA Clean & Concentrator-5 kit (Zymo)***. Details are as follows.

37.1. Add 35 ul nuclease-free water to bring the volume of sample to 50 ul.

37.2. Add 100 uL RNA Binding Buffer to each sample and mix.

37.3. Add 150 uL of ethanol (95-100%) and mix.

37.4. Transfer the sample to the Zymo-Spin™ IC Column in a Collection Tube and centrifuge 1 min with soft acceleration. Put the flow-through on the column again and centrifuge for 1 min. Discard the flow-through.

37.5. Add 400 µl RNA Prep Buffer to the column and centrifuge for 30 sec. Discard the flow-through.

37.6. Wash twice with RNA Wash Buffer: first wash: 700 µl, 30 sec; second wash: 400 ul, 2 min. Discard the flow-through.

37.7. Transfer the column in a RNase-free tube.

37.8. Elute with 20 ul DNase/RNase-Free Water directly to the column matrix. Centrifuge, take the flowthrough and apply it to the column again.

*At this stage the eluted RNA can be used immediately or stored at -80°C.

LIGATION PREPARATION

38. Prepare the T4 PNK reaction:

38.1. In a PCR tube, mix:

| **Component** | **Volume/reaction** |
|---|---|
| Eluted RNA | 18 ul |
| 10 X T4 PNK buffer | 2.2 ul |
| Superase-In 20 U/ul | 1 ul |
| T4 PNK, 10 U/ul | 1 ul |
| Total volume | 22.2 ul |

38.2. Incubate for 30 min at 37 °C.

38.3. Add 1 ul 10 mM ATP and incubate for a further 30 min at 37 °C.

39. Follow the RNeasy MinElute Cleanup Kit with the following modifications:

39.1. Add 27 ul RNase-free water, 350 ul RLT buffer, and 600 uL 100% ethanol and mix.

39.2. Transfer the sample to an RNeasy MinElute spin column and centrifuge for 15 sec and discard the flow-through.

39.3. Wash with 500 ul Buffer RPE and centrifuge for 15 sec. Discard the flow-through.

39.4. Wash with 500 ul 80% ethanol and centrifuge for 2 min. Discard the flow-through.

39.5. Transfer the column into a new 2 ml collection tube and centrifuge full speed for 5 min.

39.6. Transfer the column in a RNase-free tube.

39.7. Elute with 10 ul DNase/RNase-Free Water directly to the column matrix. Centrifuge, take the flowthrough and apply it to the column again.

SEQUENCING LIBRARY CONSTRUCTION

40. Follow ***NEBNext Small RNA Library Prep Set for Illumina*** protocol. Briefly:

40.1. In a RNase-free PCR tube, mix:

- 6 ul RNA sample
- 1 ul green 3’ SR Adaptor

40.2. Put immediately on ice.

40.3. Set the thermocycler at 25°C.

40.4. Add:

- 10 ul 3’ Ligation Reaction Buffer (pipette slowly due to the solution’s viscosity)
- 3 ul 3’ Ligation Enzyme Mix

40.5. Incubate 1 hour at 25°C.

40.6. Add:

- 4.5 ul nuclease-free water
- 1 ul SRRT Primer

40.7. Incubate following NEBNext protocol in the thermocycler.

40.8. Ligate 5’ SR Adaptor (skip steps 7-8 of the NEBNext protocol)

40.9. Reverse transcription (see NEBNext protocol) - Set lid = 100C.

40.10. PCR (see NEBNext protocol WITH THE FOLLOWING MODIFICATION)

*** Use custom illumina multiplexing primer instead of index 1 primer at the amplification step. Also, reduce the volume of the SR Primer for Illumina and the Index Primer from 2.5 μL each to 2 μL each.** So, add:

- 50 ul LongAmp Taq2X Master Mix
- 2 ul SR Primer
- 2 ul Illumina TrueSeq Adaptor 25 uM
- 1 ul SyBR dye
- 5 ul RNase-DNase-free water

40.11. Perform the PCR amplification in 2 qPCR tubes (50 ul/tube).

40.12. Set “Quick Plate 96 wells SYBR”

40.13. Follow amplification on real time: stop when the amplification reaches the beginning of the plateau (at the next 70°C step)

PURIFY QPCR PRODUCTS

41. Follow the ***Select-a-size DNA Clear & Concentrator protocol (Zymo)***, with:

  - cutoff = 100 bp. (i.e. ethanol volume = 100 ul).
  - Elution volume = 15 -20 ul

MEASURE DNA CONCENTRATION

42. Follow Qubit DNA High Sensitivity kit.

Briefly, mix in a Qubit tube:

- 199 ul Qubit HS Buffer
- 1 ul Qubit HS Reagent
- 1 ul of DNA sample

And measure the concentration on the Qubit 2.0 fluorometer.

**Purified PCR products can be stored at 4°C.**

CHECK DNA FRAGMENTS SIZE

43. In a TapeStation Optical Tube, mix:

  - 0.5 ul DNA sample
  - 1.5 ul water
  - 2 ul HS Sample Buffer

44. Place the tube, the tape and the TapeStation tips on the TapeStation 4200 and follow the steps on the computer (use Electronic Ladder).

45. Measure the average size of the fragments (create a region around the most important peak).

46. Save report.

*NB: Appropriate fragments size should be ~165 bp.*
*If not, proceed to size-selection of libraries.*

RECIPE 1 Lysis Buffer

| Components | Volume | Volume | Volume |
|---|---|---|---|
| Tris pH 8.0, 1 M | 75 ul | 100 ul | 175 ul |
| $NH_{4^{\circ}C}l$, 1 M | 75 ul | 100 ul | 175 ul |
| MgOAc, 1 M | 30 ul | 40 ul | 70 ul |
| Superase-In, 20 U/µL | 45 ul | 60 ul | 105 ul |
| 5′-guanylyl imidodiphosphate (GMPPNP), 10 mg/mL | 3 ul | 4 ul | 7 ul |
| Chloramphenicol (CAM), 50 mg/mL | 30 ul | 40 ul | 70 ul |
| RNase-free water | q.s. | q.s. | q.s. |
| **Total Volume** | **3 mL** | **4 mL** | **7 mL** |

RECIPE 3 MNase Reaction Mix

| Components | Volume |
|---|---|
| Clarified Lysate, 25 $A_{260}$ Units | 196 - X (*) ul |
| $CaCl_2$, 500 mM | 2 ul |
| Superase-IN, 20 U/ul | 2 ul |
| NEB MNase 500 U/ul | X (*) ul |
| **Total Volume** | **200 ul** |

***(*) to be calculated according to OD measurements (see protocol step 11)***

RECIPE 4 Polysome Buffer (10 mL) - **1.5 x 2 x number of samples - to be stored on ice**

| Components | Volume | Volume | Volume |
|---|---|---|---|
| $MgCl_2$, 1 M | 500 ul | 750 ul | 2 mL |
| EGTA, 0.5 M | 500 ul | 750 ul | 2 mL |
| Tris-HCl pH 8.0, 1 M | 500 ul | 750 ul | 2 mL |
| NaCl, 5 M | 500 ul | 750 ul | 2 mL |
| Nonidet P40 (NP40), 100% | 100 ul | 150 ul | 400 ul |
| RNase-free water | q.s. | q.s. | q.s. |
| **Total Volume** | **10 mL** | **15 mL** | **40 mL** |
